# The *Botrytis cinerea* Crh transglycosylae is a cytoplasmic effector triggering plant cell death and defense response

**DOI:** 10.1101/2020.06.23.166843

**Authors:** Kai Bi, Loredana Scalschi, Gupta Namrata Jaiswal, Renana Frid, Wenjun Zhu, Gal Masrati, Tesfaye Mengiste, Amir Sharon

## Abstract

Crh proteins catalyze crosslinking of chitin and glucan polymers in the fugal cell wall. We revealed a novel and unexpected role of *Botrytis cinerea* BcCrh1 as a cytoplasmic effector and elicitor of plant defense. During saprophytic growth the BcCrh1 protein is localized in vacuoles and ER. Upon plant infection the protein accumulates to high levels in infection cushions, it is then secreted to the apoplast and translocated into plant cells, where it induces cell death and defense responses. Two regions of 53 and 35 amino acids were found sufficient for protein uptake and cell death induction, respectively. Dimerization of BcCrh proteins was necessary for the transglycosylation activity and proper fungal development, while the monomeric proteins was sufficient for induction of cell death. *Arabidopsis* lines expressing the *bccrh1* gene had reduced sensitivity to *B. cinerea,* demonstrating the potential use of the protein in plant immunization against necrotrophic pathogens.

## Introduction

*Botrytis cinerea* is a wide host-range necrotrophic plant pathogen causing severe global damages^1, 2^,. The infection process includes an early stage, characterized by local necrotic lesions, an intermediate stage during which the lesion begins to spread at increasing rate, and a late stage of constant lesion spreading^3^. A working model derived from these findings predicts secretion of cell death-inducing factors during the early stage that promote formation of patches of dead tissue, which serve as foci for the following stages^4^. The model has been supported by the discovery of secreted proteins with cell death-inducing activity that are collectively referred to as necrosis-inducing proteins (NIPs)^5^. Broadly, all NIPs can be divided into two main classes: proteins that lack a known domain, and secreted enzymes that also induce plant cell death (will be referred to as catalytic NIPs).

The best studied NIPs are small secreted proteins that were collectively named NEP or NELP/NLP (NEP-like proteins)^6, 7^. Proteins in this family induce hypersensitive-like cell death in a variety of dicotyledonous but not in monocotyledonous plants^8, 9, 10^. Catalytic NIPs are all glycosyl hydrolases (GHs) that degrade plant cell wall sugar polymers such as pectin^11^, hemicellulose^5, 12, 13^ and glucose polymers^14^, All of the GH NIPs that have been characterized so far remain in the apoplast after secretion by the fungus, and their cell death-inducing activity is mediated by plant extracellular membrane components, commonly in SOBIR-BAK1-depedent manner^5, 12, 13^. Similar to the non-catalytic NIPs, all of the tested hydrolase NIPs induce plant defense, which in many cases was found unrelated to their catalytic activity^5, 12, 13, 14^. In a few cases, a short NIP-derived peptide fragment was found sufficient to induce necrosis and activate defense^13, 15, 16^, while in other cases disruption of the tertiary structure of the protein eliminated the necrosis- and defense-inducing activities^5, 17^.

In search of novel NIPs, we have analyzed secretome from bean leaves after infection with *B. cinerea* spore suspension^5^. Here we report on the identification and characterization of BcCrh1, a GH16 transglycosidae that induces cell death and defense responses in plants. Crh (Congo red hypersensitivity) are a highly conserved family of proteins that facilitate cross-linking between chitin and glucan polymers in the fungal cell wall^18, 19, 20^. Hence, unlike other catalytic NIPs, BcCrh1 is not involved in plant cell wall degradation, but rather in fungal cell wall biosynthesis. Furthermore, we show that the BcCrh1 protein is internalized into the plant cell and that this internalization is required for induction of cell death, meaning that BcCrh1 is a cytoplasmic effector, unlike most other NIPs that are apoplastic. We also found that BcCrh1 forms homodimers, as well as heterodimers with other Crh members, which is necessary for the fungal cell wall biosynthesis, while the monomeric protein is involved in fungal-plant interaction. Collectively, our study reveals a novel and unexpected role of Crh proteins as mediators of fungal-plant interaction, and provides new details on their role in cell fungal wall biosynthesis.

## Results

Proteomic analysis of *B. cinerea* secretome that was collected from infected leaves 28 hpi revealed presence of 259 predicted proteins^5^. In search for cell death inducing proteins, we cloned selected candidate genes and transiently expressed them in *Nicotiana benthamiana* leaves using *Agrobacterium tumefaciens*-mediated transformation (agroinfiltration assay). Bcin01g06010, which induced strong cell death in *N. benthamiana* was further characterized.

### Bcin01g06010 is a Crh family protein

Sequence analysis of the Bcin01g06010 protein using SignalP-5.0 (http://www.cbs.dtu.dk/services/SignalP/) and TMHMM version 2.0 (http://www.cbs.dtu.dk/services/TMHMM/), predicted a secretion signal at the N-terminal end and absence of transmembrane helices, indicating that Bcin01g06010 is a secreted protein, in accordance with its presence in *B. cinerea* secretomes^5, 21, 22^. SMART (http://smart.embl.de/) analysis showed presence of a conserved glycosyl hydrolase family 16 (GH16) domain between amino acid residues 49-263 (E-value 1.5e-36). BLAST searches revealed similarity to proteins in the Crh family (Supplementary Fig. 1), and 3D structure prediction showed strong structural similarity (Probability 100%, E-value 1.1e-32) to *A. fumigatus* AfCrh5 (Supplementary Fig. 2a). The protein was named BcCrh1 based on the homology to the *Saccharomyces cerevisiae* Crh1 protein. BLAST search of the NCBI database and published RNA sequencing data^23^ revealed three additional Crh proteins members that were named BcCrh2 (Bcin15g03070), BcCrh3 (Bcin13g03640) and BcCrh4 (Bcin07g04870). All of the four *B. cinerea* Crh proteins had a predicted secretion signal and conserved a GH16 domain (Supplementary Fig. 1a). Sequence alignment of the *B. cinerea* Crh family proteins with *S. cerevisiae*, *C. albicans* and *A. fumigatus* Crh homologues revealed the DEXDXE motif (enzymatic activity site) and the GTIXWXGG motif (the acceptor sugar binding site), that are highly conserved in all Crh proteins (Supplementary Fig. 1b, c).

### The BcCrh1 peptide is a cell death–inducing cytoplasmic effector

BcCrh1 triggered local cell death in *N. benthamiana* and tomato leaves within five and two days following Agrobacterium infiltration or treatment with purified protein solution, respectively (Supplementary Fig. 3). To test if the glycosyl hydrolase enzymatic activity of BcCrh1 was necessary for cell death-inducing activity, we mutated the conserved catalytic residues (E120Q/D122H/E124Q) to generate the MBcCrh1 enzyme inactive protein. Infiltration of leaves with an Agrobacterium strain harboring 35S:MBcCrh1 vector, or with purified MBcCrh1 protein, produced similar necrotic spots as infiltration with the native BcCrh1 protein (Supplementary Fig. 3a, b), which indicated that induction of plant cell death is unrelated to the enzymatic activity of BcCrh1. Expression of the proteins was verified by western blot analysis (Supplementary Fig. 4). Similar results were obtained from protein-infiltration assay of *N. benthamiana* leaves with purified BcCrh1 and MBcCrh1 proteins at concentrations ranging from 55 to 2.75 μM (Supplementary Fig. 5a). No necrosis was induced in *Arabidopsis* (Supplementary Fig. 5b), and similar to other *B. cinerea* NIPs, BcCrh1 did not promote necrosis in monocots (Supplementary Fig. 5c).

To determine the site of interaction with the plant, we performed agroinfiltration assay of *N. benthamiana* leaves with two types of constructs for expression of the native and enzymatic inactive proteins: plasmids BcCrh1^21–391^ and MBcCrh1^21–391^, in which the N-terminal secretion sequence was deleted, and plasmids SP(PR3)-BcCrh1^21–391^ and SP(PR3)-MBcCrh1^21–391^ in which the native secretion signal was replaced with the secretion sequence of *Arabidopsis* pathogenesis-related protein 3 (PR3). The first set of constructs will result in intracellular protein accumulation; the second set will result in secretion of the protein outside of the plant cell. Transfection of *N. benthamiana* leaves with Agrobacterium expressing any of the four constructs induced similar levels of cell death (Fig. 1a, Supplementary Fig. 4). These results indicate that cytoplasmic localized BcCrh1 or MBcCrh1 induced plant cell death. To further investigate the localization of secreted BcCrh1 in plant tissues, we infiltrated plant leaves with Agrobacterium expressing a BcCrh1/MBcCrh1-GFP fusion protein with a secretion signal. Confocal fluorescence microscopy showed that the fusion protein accumulated inside the plant cells, unlike control leaves that were treated with GFP-expressing Agrobacterium, in which the fluorescent signal was observed exclusively in the apoplastic space (Fig. 1b, Supplementary Fig. 4). These results confirmed the cytosolic localization of BcCrh1.

**Figure 1.**
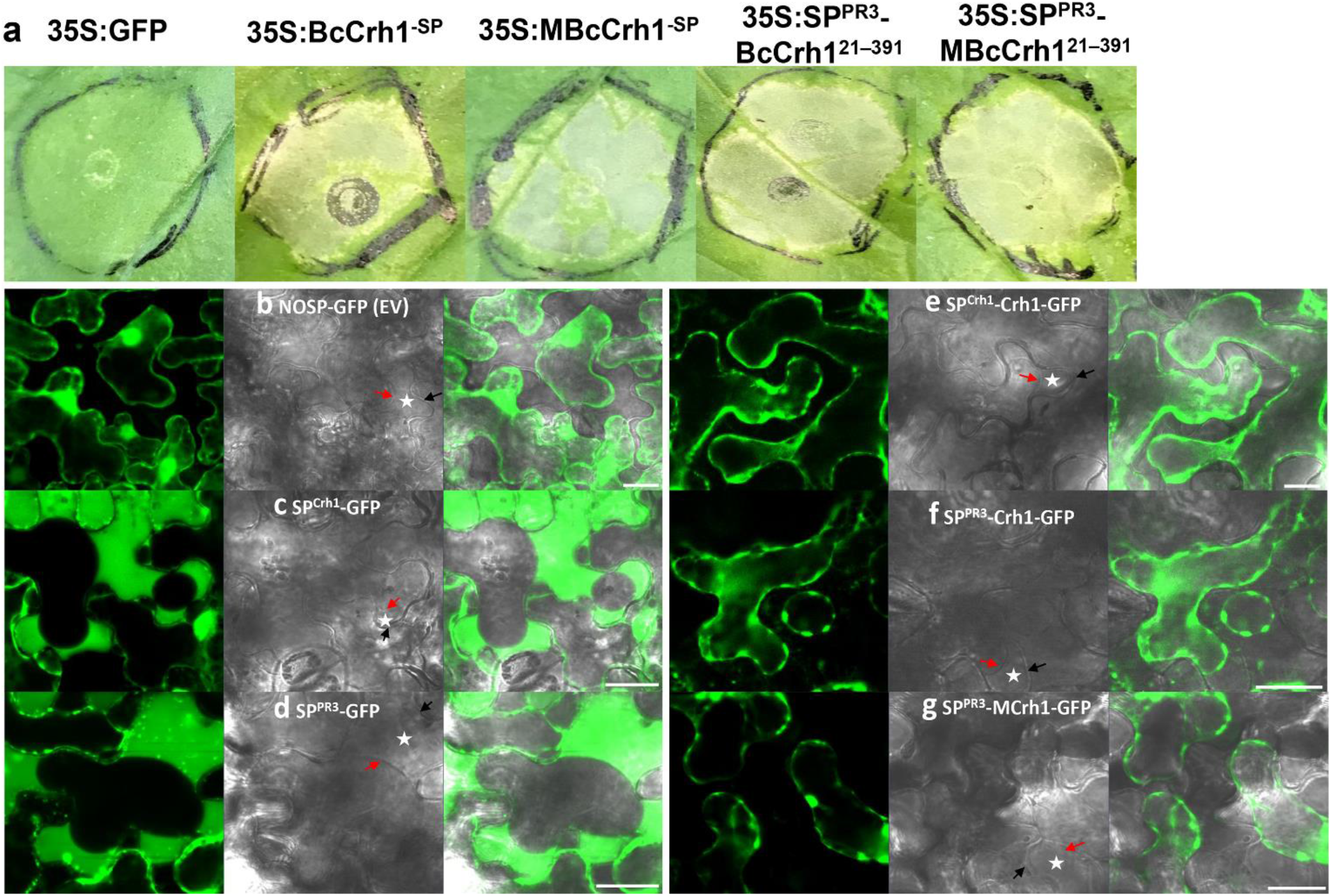
Localization of BcCrh1 inside plant cells is required for cell death induction. **a**, Plants were infiltrated with Agrobacterium strains that were transformed with the *bccrh1* gene with or without secretion signal (SP). Images of necrotic lesions (**a**) were taken five days after agroinfiltration. 35S:GFP – free GFP (control). 35S:BcCrh1^−SP^ and 35S:MBcCrh1^−SP^: native and enzyme inactive BcCrh1 respectively, without secretion signal, 35S:SP^PR3^-BcCrh1^21–391^ and 35S:SP^PR3^-MBcCrh1^21–391^: fusion of the native and enzyme inactive BcCrh1 respectively, with plant functional signal peptide PR3. **b-g**, Subcellular localization of GFP-fusion proteins. Leaves were harvested two days after agroinfiltration, submerged for 20 min in 0.8 M mannitol to induce plasmolysis, and then samples were analyzed by a confocal microscope. Typical apoplastic space between the cell wall (black arrow) and plasma membrane (red arrow) in plasmolysed plant cells is marked with white asterisks. Left panel shows images of cells following infiltration with GFP (control) constructs, right panel shows images following infiltration with BcCrh1-GFP fusion constructs. (**b**) - GFP without SP, (**c**) - GFP fused to the BcCrh1 SP, (**d**) - GFP fused to PR3 SP, (**e**) - BcCrh1-GFP with native SP, (**f**) - BcCrh1-GFP fused to PR3 SP, (**g**) - MBcCrh1-GFP fused to PR3 SP. Bars = 20 μm.

### Translocation to host cells and induction of cell death are mediated by specific regions of BcCrh1

In search of specific parts of BcCrh1 that might mediate protein uptake by plant cells, we performed agroinfiltration assays of *N. benthamiana* leaves with constructs that contained the secretion sequence and various segments of the protein. The expectation was that the protein segments that contain the uptake signal will be first secreted out of the plant cell and then induce cell death following processing of the secretion signal and uptake by the plant cells (as in Fig. 1), while no necrosis is expected by segments that lack the uptake signal. These analyses revealed that deletion of amino acids 21-74 (BcCrh1^21-74^) prevented induction of necrosis (Fig. 2), suggesting that the first 53 aa that follow the secretion signal mediate uptake of the BcCrh1 protein by plant cells. To verify that BcCrh1^21-74^ contains a functional uptake signal, we fused the BcCrh1^1-74^ (includes the secretion signal and the putative uptake signal) to the apoplastic NIP BcXYG1^5^. Agroinfiltration with this construct failed to induce cell death, unlike control treatment with the native BcXYG1, which induced strong necrosis at 3 dpi (Fig. 3a, Supplementary Fig. 4). Consistent with the agroinfiltration result, infiltration of leaves with purified BcCrh1^21-144^ peptide caused significant cell death at 2 dpi, while treatment of leaves with the BcCrh1^75-144^ peptide that lacks the putative uptake signal caused no visible symptoms (Fig. 3b). In order to obtain more direct evidence for the translocation function of the 53 aa region, we generated GFP fusion with the BcCrh1^21-144^ and BcCrh1^75-144^ peptides and administered the purified proteins to a tomato cell suspension-culture. After 18 h of incubation, GFP signal was detected inside tomato cells that were incubated with the GFP-BcCrh1^21-144^ protein, while in cells that were incubated with the GFP-BcCrh1^75-144^ protein there was no signal in the cytoplasmic space (Fig. 3c). Taken together, these results confirm that BcCrh1 is an intracellular, cell death-inducing effector, and that translocation into the plant cells is mediated by the N’ 53 amino acids residues of the protein.

**Figure 2.**
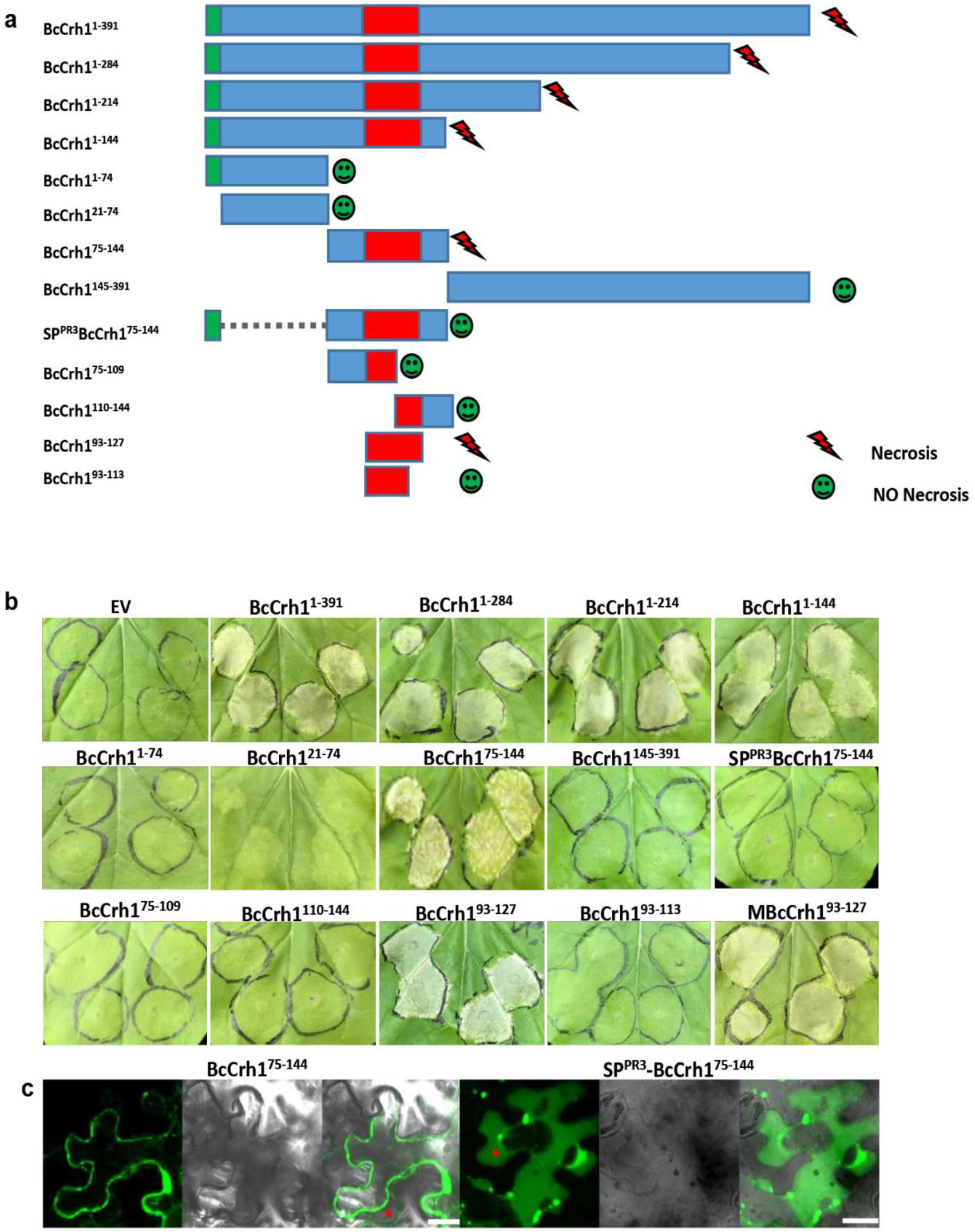
A peptide of 35 amino derived from BcCrh1 is necessary and sufficient for induction of plant cell death. **a**, Schematic presentation of different BcCrh1 fragments that were constructed and used in agroinfiltration assay. Green – SP, Red – the minimal region that was found sufficient for induction of cell death, Bleu – other regions of the tested fragment. **b, c**, Plants were infiltrated with Agrobacterium strains that contain constructs encoding the different BcCrh1 fragments (**a**). **b**, Pictures were taken five days after leaf infiltration. **c**, Two days after agroinfiltration leaves were submerged for 20 minutes in 0.8 M mannitol to induce plasmolysis and then samples were analyzed by a confocal microscope. Typical apoplastic space between the cell wall and plasma membrane in plasmolysed plant cells is marked with red asterisks. Bars = 20 μm.

**Figure 3.**
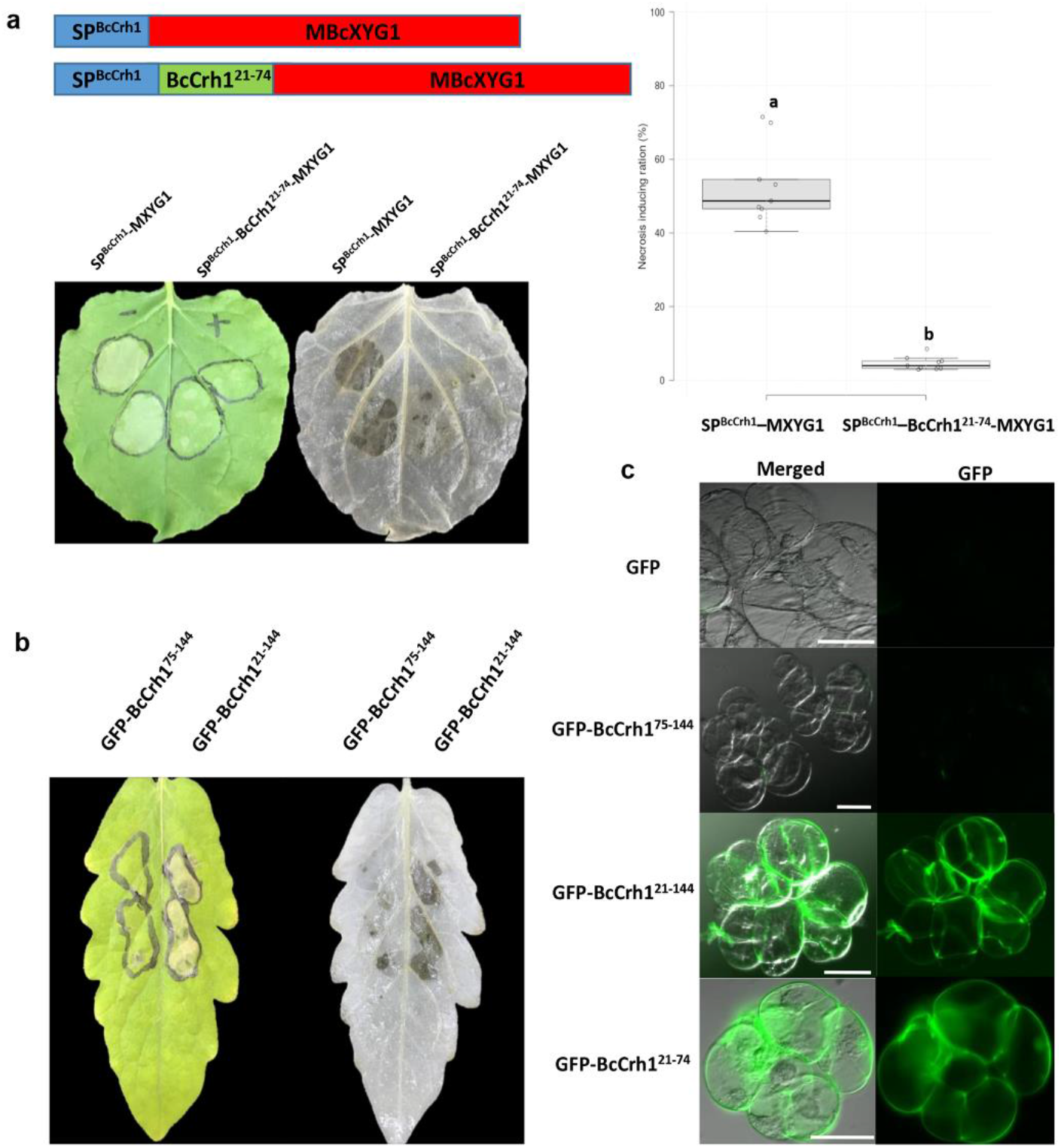
Uptake of BcCrh1 by plant cells is mediated by 53 amino acids located at the N’ end of the protein. Following analysis of a series of deletion constructs, a minimal region suspected as necessary for protein uptake was defined. **a**, To test the delivery of heterologous proteins using this signal, we generated two expression constructs with the apoplastic NIP BcXYG1 (Zhu et al., 2017): fusion of the BcCrh1 SP (blue) to MBcXYG1 (red), and fusion of the BcCrh1 SP together with the suspected delivery sequence (green) to MBcXYG1. *N. benthamiana* leaves were agroinfiltrated with the different constructs and pictures were taken after three days. The SP^BcCrh1^-MBcXYG1 fusion construct (which is apoplastic) caused severe necrosis, while induction of necrosis by the MBcXYG1 fused to the SP with uptake signal is compromised. The agroinfiltrated leaves were bleached with ethanol and the ratio of necrotic area (necrosis/infiltrated area) was calculated. Center lines of the Boxplots show the medians, box limits indicate the 25th and 75th percentiles; whiskers extend 1.5 times the interquartile range from the 25th and 75th percentiles; all present data are plotted as open circles. Nine sample points from three independent biological replicates were used for statistical analysis. Different letters indicate statistical differences at *P* ≤ 0.01 using one-way ANOVA. **b**, Tomato leaves were infiltrated with 11 μM of purified fragments of the BcCrh1 protein: BcCrh1^21-144^ and BcCrh1^75-144^ and pictures were taken after 48 h. Leaves that were infiltrated with the BcCrh1^21-144^ fragment that contains the putative uptake and necrosis inducing domains developed strong necrosis, whereas the BcCrh1^75-144^ couldn’t induce necrosis symptoms. **c**, Tomato cell cultures (MsK8) were incubated with 5.5 μM of GFP-tagged peptide solutions (GFP-BcCrh1^21-144^ and GFP-BcCrh1^75-144^) for 18 h, then subsequently washed three times in incubation buffer and visualized by confocal laser microscopy. Bars = 50 μm.

To define the minimal region required for cell death-inducing active of BcCrh1, we generated a series of N-terminal and C-terminal deletions and tested their cell death-inducing activity. Long deletions at the C’ end including BcCrh1^1-284^, BcCrh1^1-214^, and BcCrh1^1-144^ retained full activity, however when residues 75-284 were deleted with (BcCrh1^1-74^) or without (BcCrh1^21-74^) the secretion signal, cell death-inducing activity was lost (Fig. 2a, b). These results indicated that BcCrh1^75-144^ is necessary for cell death induction. Transient expression of BcCrh1^75-144^ but not BcCrh1^145-391^, induced cell death in *N. benthamiana* leaves, confirming that the region between amino acids 75-144 was both necessary and sufficient for induction of plant cell death. When BcCrh1^75-144^ was fused to the PR3 secretion signal (SP^PR3^-BcCrh1^75-144^), the cell death activity was lost (Fig. 2a, b). Confocal images of *N. benthamiana* cells transiently expressing BcCrh1^75-144^ or SP^PR3^-BcCrh1^75-144^ GFP fusions, confirmed intracellular and apoplastic localization of the proteins, respectively (Fig. 2c). Following further analyses of different parts of the protein, the region necessary for induction of cell death was narrowed down to 35aa between residues 93-127 (Fig. 2a, b). Because BcCrh1^93-127^ contains the catalytic pocket, we mutated the catalytic residues (E120Q/D122H/E124Q) in BcCrh1^93-127^. The mutated epitope (MBcCrh1^93-127^) still triggered cell death (Fig. 2b), verifying the earlier results that were obtained with MBcCrh1.

### BcCrh1 Triggers PTI and induces host resistance against *B. cinerea*

Infiltration of *N. benthamiana* and tomato leaves with Agrobacterium or purified protein solution resulted in accumulation of reactive oxygen species (ROS) and activation PTI genes, respectively (Fig. 4a, b). When plants were inoculated with *B. cinerea* 48 h after infiltration with the protein, infection was significantly reduced compared to control plants that were pre-treated by infiltration of GFP protein (Fig. 4c). Thus, along with cell death inducing activity BcCrh1 also triggers plant defense responses, similar to the vast majority of *B. cinerea* NIPs^5, 12, 13, 14, 15, 16, 24, 25^. Since BcCrh1 does not induce cell death in *Arabidopsis thaliana*, we produced *A. thaliana* transgenic lines that express the BcCrh1 protein and tested their sensitivity to infection by the fungus (Supplementary Fig. 4). The BcCrh1 expressing plants were much less sensitive to infection by *B. cinerea* compared with control plants that were transformed with an empty vector, as determined by lesion size 3 dpi (Fig. 4d). These results show that BcCrh1 induces PTI reaction, which can be used to improve plant resistance to *B. cinerea*.

**Figure 4.**
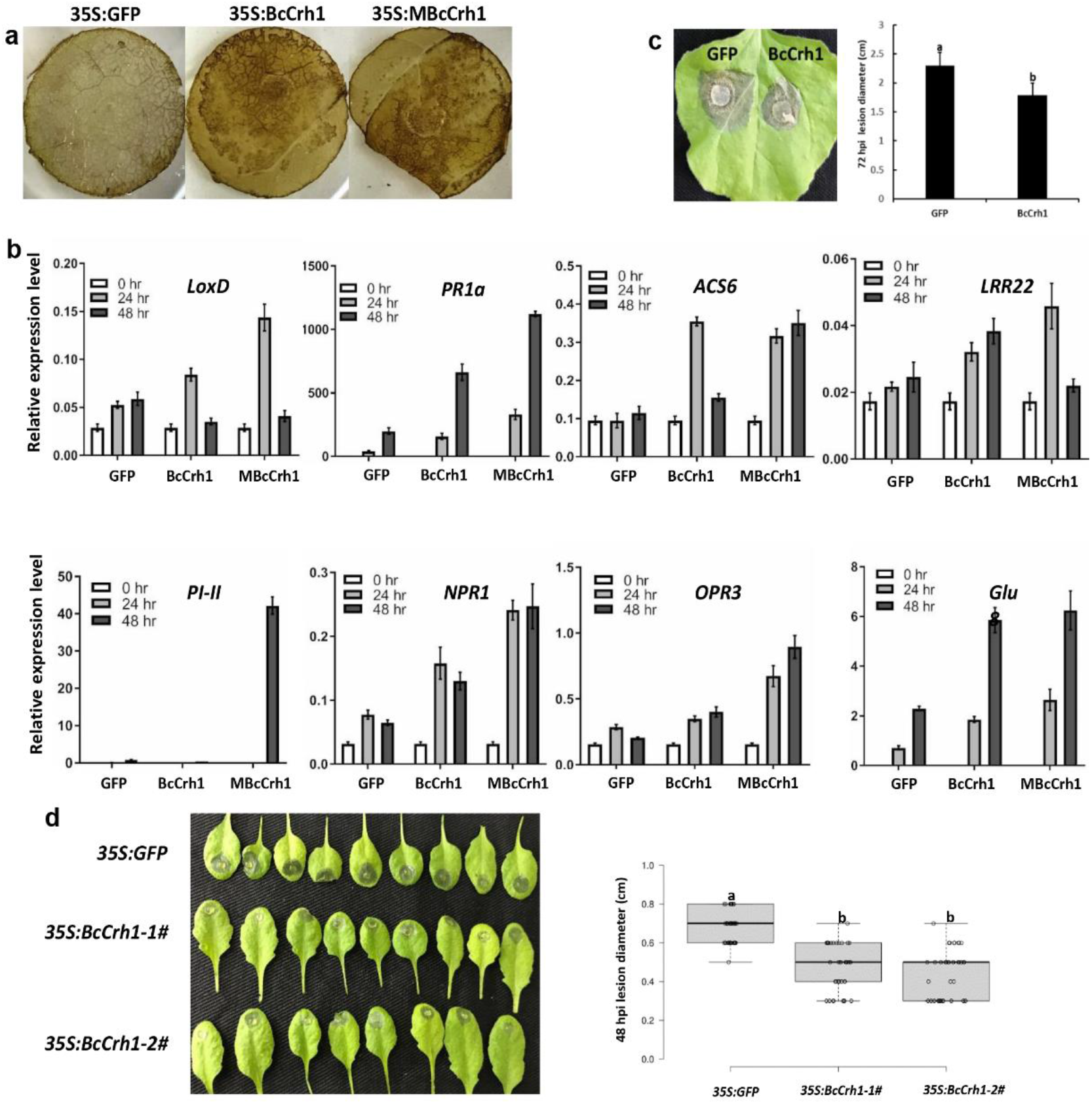
BcCrh1 triggers plant immunity responses and enhances plant resistance to Botrytis infection. **a**, Accumulation of reactive oxygen species (ROS) in *N. benthamiana* two days after agroinfiltration with construct encoding GFP (35S:GFP), and native (35S:BcCrh1) or enzymatic inactive (35S:MBcCrh1) BcCrh1. Leaves were stained with DAB and then decolorized with ethanol to allow easier visualization of reduced DAB precipitate. **b**, Relative expression of tomato defense genes 24 and 48 h after infiltration with GFP, BcCrh1 and MBcCrh1. Gene expression levels were determined by qRT-PCR and normalized using the transcription level of the *actin* gene. Values represent mean ± sd (n = 9) from three independent biological replicates and three technical replicates. **c**, Infection assay of *N. benthamiana* leaves after infiltration with purified proteins. Following leaf treatment, the plants were kept in a growth chamber for 48 h and then inoculated with mycelium-agar of *B. cinerea*. The plants were incubated in a moist chamber and symptoms were recorded 72 hpi. Data represent means ± sd from three independent experiments, each with four replications. **d**, Infection assay of *A. thaliana* GFP (*35S:GFP*) and BcCrh1 transgenic plants (*35S:BcCrh-1# and 35S:BcCrh-2#*). Leaves were inoculated with droplets of *B. cinerea* spore suspension, incubated in a moist chamber and symptoms were recorded 48 hpi. Center lines of Boxplots show the median values; box limits indicate the 25th and 75th percentiles; whiskers extend to minimum and maximum values; data points are plotted as open circles. n = 44, 36, 36 sample points from three independent biological replicates. Different letters indicate statistical differences at P ≤ 0.01 using one-way ANOVA.

### BcCrh1 is expressed in planta and secreted from infection cushions

Crh proteins are fungal cell wall transglycosidaes^18, 19, 20^. Nevertheless, BcCrh1 was found in the fungal secretome, it is internalized by the plant cells, where it triggers plant cell death and defense responses, implying that it also affects fungal-plant interaction. To gain insight on possible role of BcCrh1 in infection development we studied *bccrh1* gene expression and protein localization in planta. Transcript levels of *bccrh1* increased sharply following infection and reached a peak 12 hpi, in contrast to much more gradual increase on solid Gamborg’s B5 medium that peaked at 60 h post germination (Supplementary Fig. 6). To determine protein localization, a transgenic *B. cinerea* strain was produced with a *bccrh1*-*gfp* fusion gene under control of the native *bccrh1* promoter. During saprophytic growth on culture medium the protein was observed in the fungal vacuoles and ER, and accumulated to high levels in infection cushions (Fig. 5a, b). Infection assay of onion epidermis cells showed that the protein was secreted from hyphal tip 12 - 21 hpi, and from infection cushions at 36 hpi (Fig. 5c). Further evidence for BcCrh1 secretion was confirmed by Z-series projection, demonstrating that the GFP-BcCrh1 fusion protein is first delivered to the hyphal tip and later concentrates in infection cushions (Supplementary Fig. 7). Following secretion of the protein from the fungal cells, the GFP signal diffused in the surrounding apoplastic space and then accumulated inside the onion cells, including neighboring cells in regions of the tissue that are adjacent to the invasion area (Fig. 5d). These results show different expression and localization pattern of BcCrh1; During saprophytic growth the gene is expressed constantly at moderate levels and the protein is localized inside the fungal cells, while in planta the gene is highly and transiently expressed following first contact of the fungus with the plant, and the protein accumulates to high levels in infection structures before being released to the plant apoplast.

**Figure 5.**
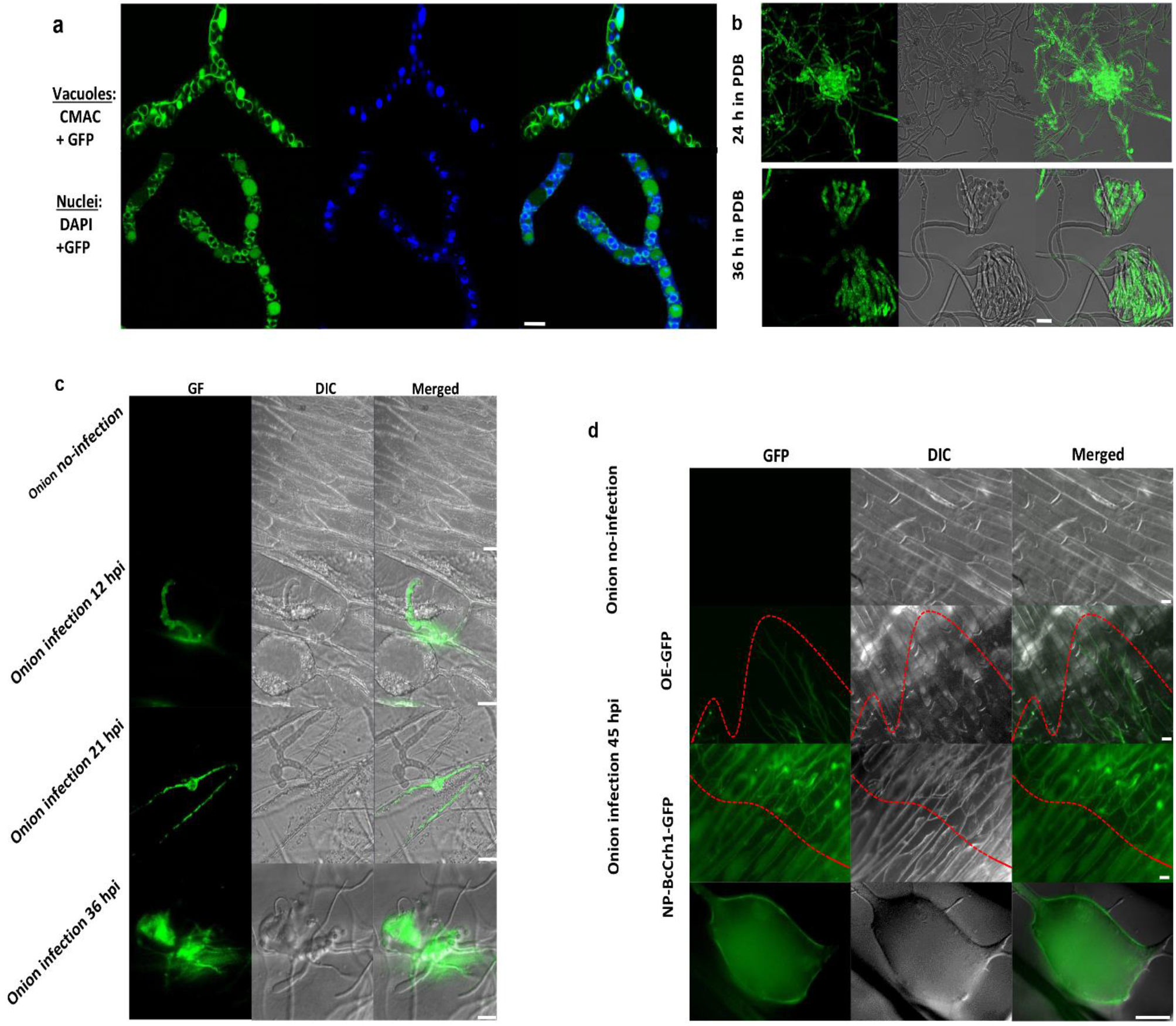
Subcellular localization of BcCrh1 during the saprophytic growth and host infection. **a**, Intracellular localization during saprophytic growth. Spores of NP-BcCrh1-GFP strain were cultured in liquid PDB medium for 12 h, the germinated spores were collected, vacuoles were stained with CMAC (top) and nuclei were stained with DAPI (bottom). Samples were visualized using a Confocal microscope. Overlap of the CMAC and GFP signals indicate vacuole localization of the BcCrh1 protein. GFP fluorescence is also observed around nuclei and other ER structures. Scale bars = 5 μm. **b**, Spores were germinated on a glass slide in PDB medium. Images were taken after incubation for 24 and 36 h. The GFP signal accumulated in the entire mycelium and in initiating infection cushions after 24 h (top), and in mature infection cushions after 36 h (bottom). Scale bar = 20 μm. **c-d**, Confocal images of onion epidermis cells that were inoculated with spores of the NP-BcCrh1-GFP strain. **c**, Images showing secretion of the protein from hyphal tips at 12, 21 hpi, and from infection cushions at 36 hpi. Scale bars = 50 μm at 12 hpi and 20 μm in all other images. **d**, Intracellular localization of secreted BcCrh1-GFP protein in plasmolysed onion cells 45 hpi. BcCrh1-derived GFP signal was significantly enriched in the cytoplasmic space of plasmolyzed onion cells adjacent to the invasion area. Onion cells without inoculation or inoculated with spores of OE-GFP strain were used as controls. The red dotted line indicates the leading edge of infection hypha. Scale bars = 20 μm.

### BcCrh1 is dispensable for *B. cinerea* pathogenicity and development

Deletion or over expression of the *bccrh1* gene had no visible effect on fungal development or pathogenicity (Fig. 6; Supplementary Fig. 8a - d). to determine if the enzymatic activity activates a defense response that counteracts the cell death-inducing activity, two additional strains were generated that overexpress the enzymatic inactive protein (MBcCrh1) in wild-type (OE-MBcCrh1) and *bccrh1* deletion (Δ/OE-MBcCrh1) genetic background. Surprisingly, both strains were less virulent than the wild type strain. In particular, pathogenicity of the double mutant was severely defected and symptoms were restricted to local lesions (Fig. 6a, b). The Δ/OE-MBcCrh1 strain also had developmental defects, including reduced sporulation and mycelium production, and abnormal spore shape (Fig. 6e, f, Fig. 7c, d), however there was no change in spore germination (Supplementary Fig. 8b). Microscopic observation of infected leaves showed weaker penetration ability of the Δ/OE-MBcCrh1 strain (Fig. 6c, d), which was associated with impaired infection cushion formation of this mutant compared with other strains (Supplementary Fig. 8d). Furthermore, pathogenicity of the Δ/OE-MBcCrh1 strain was at least partially restored by mechanical injury of leaves prior to infection (Supplementary Fig. 8e), suggesting that the defects were specific to the penetration stage. These phenotypic changes were all more severe when the *bccrh1* gene was deleted, suggesting that they were caused by accumulation of high levels of the MBcCrh1 protein. To test this hypothesis, we generated a strain that expresses the MBcCrh1 protein under control of the native *bccrh1* promoter in the background of Δ*bccrh1* (strain Δ/NP-MBcCrh1). This strain did not show any developmental defects and it formed normal infection cushions and disease symptoms (Fig. 7), confirming that the developmental defects of the Δ/OE-MBcCrh1 strain resulted from high level accumulation of enzyme inactive protein. The results also showed that the pathogenicity defects were caused at least in part by defects in plant invasion due to deformed infection cushions. Based on these results, we predicted that BcCrh1 form protein dimers that are necessary for the transglycosylation activity.

**Figure 6.**
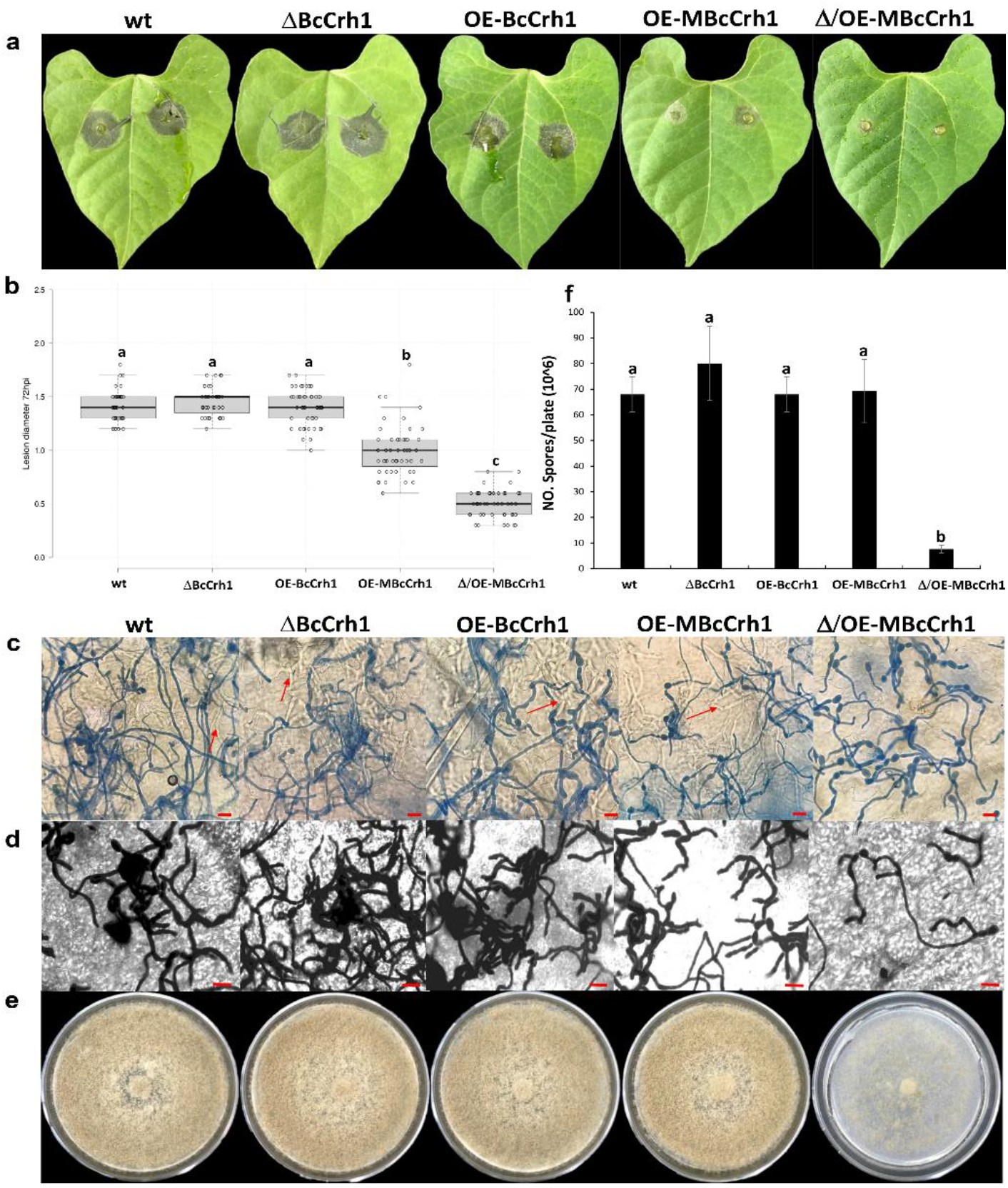
Over expression of the enzyme inactive MBcCrh1 causes reduced pathogenicity and developmental defects. **a**, Bean leaves were inoculated with spore suspensions of *B. cinerea* wild type (wt), *bccrh1* deletion (ΔBcCrh1), *bccrh1* over expression (OE-BcCrh1) and strains over-expressing the enzyme inactive (MBcCrh1) protein in wild type (OE-MBcCrh1) or *bccrh1* deletion (Δ/OE-MBcCrh1) genetic background. Pictures of symptoms were recorded 72 hpi. **b**, Average lesion diameter 72hpi. Box limits shows the 25th and 75th percentiles. The center lines of Boxplots indicate the medians values; whiskers extend 1.5 times the interquartile range from the 25th and 75th percentiles; all present data are plotted as open circles. At least 41 sample points from three independent biological replicates used for statistical analysis. **c-d**, Lactophenol cotton blue and lactophenol trypan blue staining of infected leaves. Leaf tissue was harvested at the designated time points and stained with lactophenol cotton blue that stains only the exposed hyphae (**c**) and with lactophenol trypan blue that stains both exposed and intracellular (red arrows) hyphae (**d**). Bar = 20 μm. Note almost lack of intracellular hyphae in the Δ/OE-MBcCrh1 mutant, which indicates penetration defects. **e-f**, Mycelium and spore production. Fungi were cultured on solid GB5-Glucose medium and grown at 20°C with continuous light. Pictures of the plates were taken (**e**) and spores were collected and counted (**f**) after eight days of incubation. Data collected from three independent biological replicates. Different letters indicate statistical differences at P ≤ 0.01 using one-way ANOVA.

**Figure 7.**
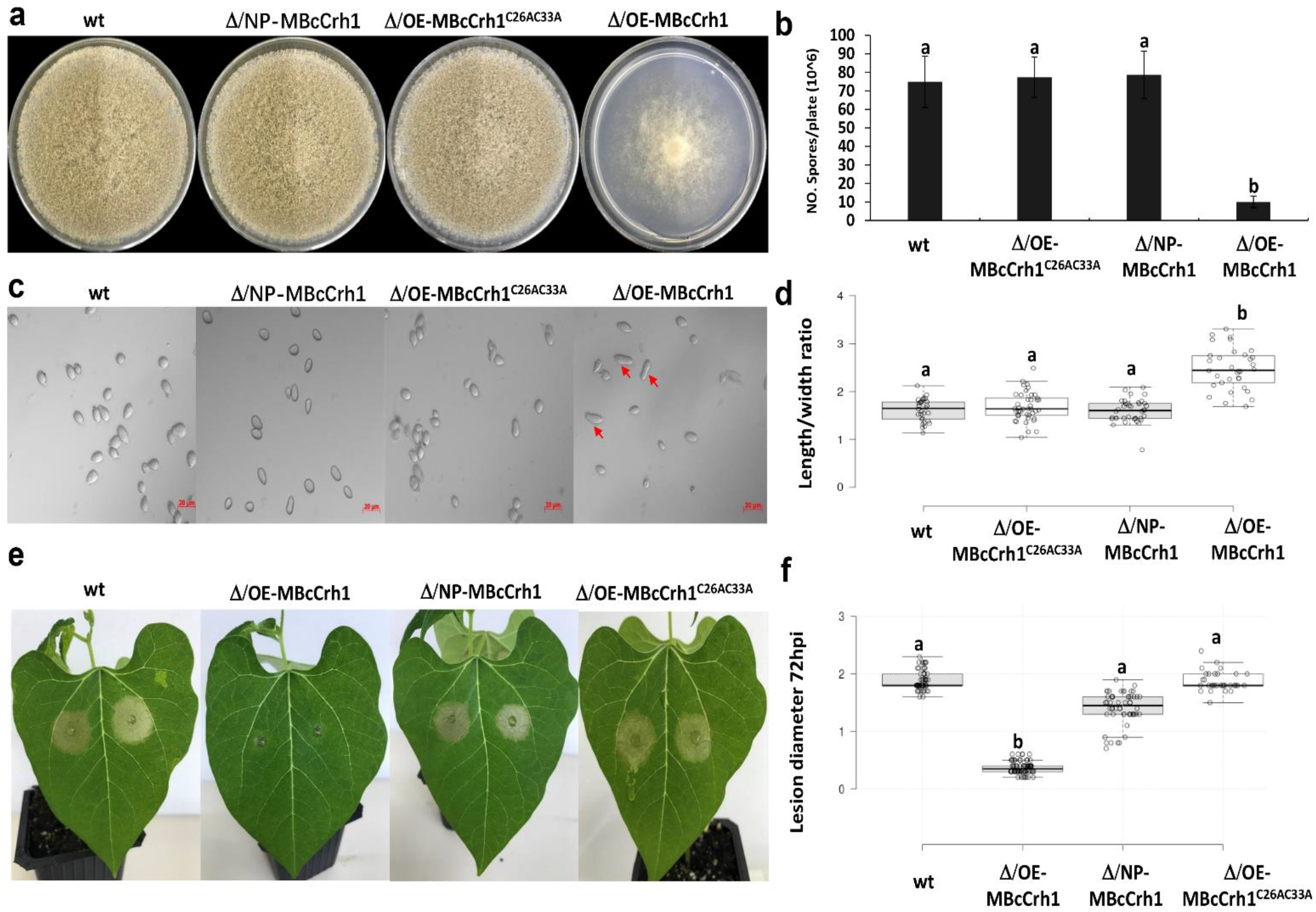
Native promoter controlled MBcCrh1 or prevention of MBcCrh1 dimerization abolish the pathogenicity and developmental defects induced by over expression of MBcCrh1. The following strains were characterized: *B. cinerea* wild type (wt), enzyme inactive BcCrh1 with the native promoter in background of *bccrh1* deletion (Δ/NP-MBcCrh1), over expression of MBcCrh1 with mutation of C26 and C33 in background of *bccrh1* deletion Δ/OE-MBcCrh1^C26AC33A^), and over expression of MBcCrh1 in background of *bccrh1* deletion (Δ/OE-MBcCrh1). **a-d**, Spore production and shape. Fungi were cultured on solid GB5-Glucose medium and grown at 20°C with continuous light. Pictures of the plates (**a**) and spores (**c**) were taken after eight days of incubation, spores were harvested and average spore numbers (**b**) and spore dimensions (**d**) were determined. Arrow indicates spores of the Δ/OE-MBcCrh1 strain with abnormal morphology. For spore numbers, data was collected from three independent biological replications. For spore dimensions, the ratio of length/width was calculated. At least 30 sample data from three independent biological replicates used for statistical analysis. One-way ANOVA was used, different letters indicate statistical differences at *P* ≤ 0.01. **e-f**, Bean leaves were inoculated with spore suspensions of the different strains, pictures (**e**) were taken and lesion size (**f**) recorded 72 hpi. Center lines of box plot show the medians, box limits indicate the 25th and 75th percentiles as determined by R software; whiskers extend 1.5 times the interquartile range from the 25th and 75th percentiles, outliers are represented by dots; data points are plotted as open circles. At least 32 sample data from three independent biological replicates used for statistical analysis. Different letters indicate statistical differences at *P* ≤ 0.01 using one-way ANOVA.

### BcCrh1 forms homodimers as well as heterodimers with other BcCrh protein members

Yeast two hybrid assay confirmed that the native as well as the enzyme inactive BcCrh1 form protein dimers (Fig. 8a). Sequence alignment of fungal Crh proteins revealed two conserved cysteine residues at positions 26 and 33 (Supplementary Fig. 1c), which form intramolecular disulfide bonds that might be important for proper folding and stability (Supplementary Fig. 2). Site directed mutagenesis of either or both C^26^ and C^33^, and deletion of the sequence including C^26-33^ showed that C^33^, but not C^26^, was required for BcCrh1 homo-dimer formation (Fig. 8c). Interestingly, despite the importance of C^33^ for homodimer formation and protein structure, the BcCrh1^C26AC33A^ mutant protein retained full necrosis-inducing activity (Fig. 8d), meaning that the cell death inducing activity is induced by the monomeric peptide and is independent of the protein tertiary structure. Yeast two hybrid assays with the three other BcCrh members showed that BcCrh2 and BcCrh3, but not BcCrh4 can also form homodimers, and that BcCrh1, BcCrh2 and BcCrh3 can form heterodimers with each other, all of which were independent of the enzymatic activity and mediated by the conserved cysteine residues (Fig. 8b, c). In order to verify our hypothesis that the excessive MBcCrh1 disrupts the normal function of the *B. cinerea* Crh proteins by formation of an enzymatic inactive protein dimers, we overexpressed the MBcCrh1^C26AC33A^, which is unable to form dimers, in background of Δ*bccrh1*. In accordance with our prediction, this strain had no growth defects and it caused normal disease symptoms (Fig. 7).

**Figure 8.**
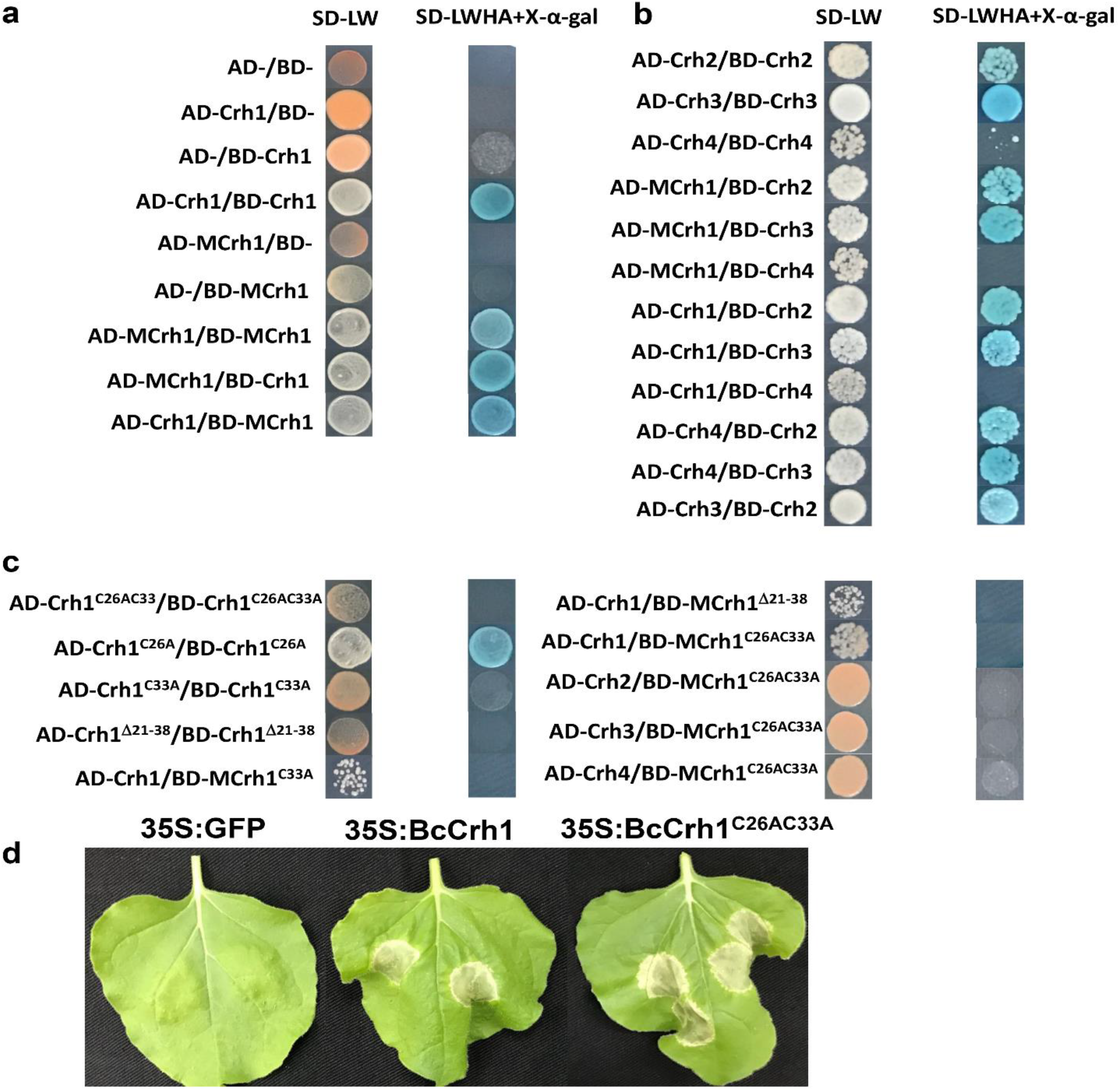
The BcCrh proteins from homo- and hetero-dimers. **a-c**, Interaction between different BcCrh members (**a, b**) and effect of mutations on the interaction (**c**) were tested by yeast 2 hybrid assay (Y2H). “-” means empty vector, “C26AC33A” indicates site-directed mutagenesis of the two cysteine residues (C26 and C33), “Δ21-38” stands for deletion of the N-terminal amino acids 21-38. SD/ Trp Leu medium was used to confirm transformation. SD/ Trp Leu His Ade medium containing X-α-gal was used for screening of yeast strains with positive protein-protein interaction. **d**, Cell death inducing activity of the BcCrh1^C26AC33A^ mutant protein was tested by agroinfiltration assay of *N. benthamiana* leaves. Pictures of representative leaves were taken five days after infiltration. The mutant protein was at least as active as the native BcCrh1 protein.

## Discussion

Glycosyl hydrolases (GH) are the largest GO group in *B. cinerea* secretome. The majority of the GHs are hydrolases of plant cell wall sugar polymers, such as cellulose, pectin, hemicellulose and other plant cell wall-associated polysaccharides^26^. Among them, a small number of proteins (catalytic NIPs) can also trigger cell death in plants^5, 12, 13, 14^, similar to the non-catalytic NIPs, such as NEP/NELP^6, 7^ and ceratoplatanins^25, 27^. All of the catalytic NIPs that have been characterized so far remain in the apoplast after secretion by the fungus, and their cell death-inducing activity is mediated by plant extracellular membrane components^5, 12, 13^. The BcCrh1 protein reported here represents a novel class of NIPs: it has a defined catalytic domain but lacks a plant cell wall degrading activity, and its site of action is inside, rather than outside the plant cell. Furthermore, the cell death-inducing activity of BcCrh1 is unexpected and surprising, since Crh family proteins catalyze crosslinking of sugar polymers in the fungal cell wall, thus it lacks clear connection to host-fungal interaction.

The Crh protein family is a conserved group of fungal-unique enzymes catalyzing the final step of cell wall maturation^18, 19, 20^. Deletion of *Saccharomyces cerevisiae crh1* or *crh2* affected cell wall integrity and proper development, and similar results were obtained in other yeast species^18, 19, 28, 29^, while deletion of any of the five Crh members in *Aspergillus fumigatus* had almost no effect on fungal development^20^. Similarly, deletion of *bccrh1* did not affect *B. cinerea* development and only slightly increased sensitivity of the fungus to congo red (Supplementary Fig. 8c). However, we found that the BcCrh proteins form dimers, and this dimerization was necessary for the transglycosylases activity. Over production of the enzyme inactive monomer resulted in severe developmental defects, indicating that two intact monomers are necessary for proper function, and further supporting a redundant role of Crh proteins in filamentous species. These results reveal a new aspect in Crh proteins function, which can lead to better understanding of their mechanistic mode of action.

Under saprophytic conditions the BcCrh1 protein was localized inside vacuoles and ER, but during pathogenic development high levels of the protein were observed in developing infection cushions, before being secreted to the plant apoplast. The fungal strains in which BcCrh dimerization was blocked had defected infection cushions and drastically reduced pathogenicity. The specific accumulation of BcCrh1 in infection cushions implies that it is necessary for infection cushion formation, which is associated with retardation of hyphal extension and massive branching. Possibly, following these pathogenicity-specific developmental events, the excessive enzyme, now serving as a virulent factor, is released from the mature infection cushions and induce plant cell death.

The BcCrh1 is a cytoplasmic effector, unlike all previously isolated catalytic NIPs, which remain in the apoplast. Delivery of the protein to the plant nucleus (by addition of NLS) prevented cell death, indicating that the site of action is in the plant cytosol (Supplementary Fig. 3c). The uptake of BcCrh1 by plant cells is pathogen-independent and mediated by the first 53 aa of the mature BcCrh1. This feature places BcCrh1 in the category of cell penetrating peptides (CPPs), a class of short peptides that facilitate cellular uptake of various molecules through endocytosis, macropinocytosis, and direct plasma membrane penetration^30^. Compared with animals, only a small number plants CPPs were reported in plants, among them several that mediate effectors uptake, mainly from oomycetes. Specific motifs that were associated with delivery of oomycetes include the RXLR motif that is common in *Phytophthora* effectors^31, 32, 33, 34^, LXLFLAK of CRN proteins, and CHCX-containing amino terminus motif in *Albugo laibachii* effectors proteins^35, 36, 37^. It has been proposed that RXLR effectors are internalized via endocytosis in a pathogen-independent manner^38, 39^, whereas the mechanisms of uptake of CRN and CHXC effectors remain elusive^33^. A relaxed RXLR-like motif has been proposed to be involved in the pathogen-independent, PI3P-mediated endocytosis of certain fungal effectors^39, 40, 41^, however to date, there is no known conserved motif that is shared by candidate cytoplasmic fungal effectors^33, 42^. Similar with other fungal cytoplasmic effectors, the BcCrh1 translocation sequence does not contain any known conserved motifs and it lacks homology to sequences that mediate translocation of effectors in other fungi. Thus, the N-terminal uptake signal of BcCrh1 may be considered a new class of CPP.

Similar to all other NIPs, along with plant cell death, BcCrh1 also elicits PTI response. The cell death and PTI have opposite effect on disease development, which together with high redundancy of NIPs precludes clear phenotype in deletion or over expression fungal strains. However, BcCrh1 did not induce cell death in *Arabidopsis,* which enabled better assessment of the induced PTI on disease development. Indeed, transgenic *Arabidopsis* plants that produce BcCrh1 were much less sensitive to *B. cinerea,* demonstrating the potential of BcCrh1, and possibly other NIPs in engineering pathogen resistance in plants.

## Methods

### Fungi, bacteria and plants

*Botrytis cinerea* B05.10 and derived transgenic strains were routinely cultured on potato dextrose agar medium (PDA, Acumedia) at 22°C under continuous fluorescent light supplemented with near UV (black) light. The transgenic fungal strains were cultured on PDA amended with 100 μg/ml hygromycin B (Calbiochem) and/or 100 μg/ml Nourseothricin (Sigma-Aldrich). *Escherichia coli* strain DH5α and BL21 (DE3) were used for plasmid construction and propagation and for protein expression, respectively. *A. tumefaciens* strain GV3101 was used for *A. tumefaciens*-mediated transient expression of proteins in plant leaves. All the bacteria were grown at 37°C on LB agar plates or in LB liquid medium supplemented with 100 μg/ml ampicillin and 50 μg/ml kanamycin. French bean (*Phaseolus vulgaris* L. genotype N9059), *N. benthamiana*, *A. thaliana* (ecotype Columbia-0), tomato (*Solanum lycopersicum*) cv. Hawaii 7998, and maize (*Zea mays*) cv. silver queen were grown in a growth room at 20°C (*A. thaliana*), or 25°C (all other plant species) with 16-h/8-h light/dark cycle.

### Bioinformatics analysis

The genomic sequence database of *B. cinerea* (https://mycocosm.jgi.doe.gov/Botci1/Botci1.home.html) was used for Blast searches of *B. cinerea* genes. The SignalP 5.0 server (http://www.cbs.dtu.dk/services/SignalP/) was used to predict the presence of signal peptides and the location of their cleavage sites in the proteins^43^. TMHMM Server v. 2.0 (http://www.cbs.dtu.dk/services/TMHMM/) was used for prediction of transmembrane helices in proteins^44^. Conserved protein domain search was performed by SMART MODE (http://smart.emblheidelberg.de/smart/change_mode.pl)^45^. The databases NCBI and UniProt were used for BLASTp searches. Multiple Sequence Alignment (MSA) was performed by Clustal Omega (https://www.ebi.ac.uk/Tools/msa/) and MView Version 1.63 was used to present the result. HHpred (https://toolkit.tuebingen.mpg.de/tools/hhpred) was used for prediction of the 3D structural models of template from PDB database. The template with the highest sequence identity was used for modeling. To produce the pairwise alignment between the two proteins, ConSurf was used to collect homologues. Sequences were collected from the clean UniProt database with 95% maximal identity between sequences and minimal 35% identity for homologues. Sequences that introduced large gaps to the alignment were discarded, and the pairwise alignment of BcCrh1 and 6IBW was deduced from the multiple sequence alignment. MODELLER (https://salilab.org/modeller/) was used to produce different models, and each model underwent a short energy minimization using GROMACS and the AMBER99SB-ILDN force field (https://www.nvidia.com/es-la/data-center/gpu-accelerated-applications/gromacs/). Finally, the model result with the predicted lowest energy was chosen. The models cover amino acids 25 to 275 (according to the amino acids numbering in UniProt entry A0A384J6C4). Each model underwent a short energy minimization using GROMACS and the AMBER99SB-ILDN force field. To test potential dimerization interfaces, PISA (https://www.ebi.ac.uk/pdbe/pisa/) and the template’s crystal structure (6IBW) were used.

### Construction of plasmid DNA

Primers used in this study are listed in Supplementary Table S1. All constructs were sequence-verified to confirm their integrity before further manipulation. Binary plasmids based on 2×35S MCS eGFP (pCNG) were used for gene expression in plants (*N. benthamiana* and *A. thaliana*). The ORF of target genes were amplified from cDNA with Phusion High-Fidelity DNA Polymerases (NEB). Signal peptide sequence deletion, C-terminus truncations, addition of nuclear localization (NLS) or myristoylation (CBL) signals, and site-directed PCR mutagenesis were conducted on the target sequences according to specific requirements. The fragments were cloned into linearized pCNG (the vector plasmid was excised with *Xba*I and *BamH*I) fused to the eGFP using an *E. coli* DH5α-mediated DNA assembly method^46^. For the generation of *B.* cinerea mutant strains, several plasmids were constructed. For deletion of the *bccrh1* gene, 500bp each of BcCrh1-5′ and BcCrh1-3′ flanking regions were amplified from B05.10 genomic DNA and added on each side of a hygromycin-resistance cassette to produce the *bccrh1* deletion plasmid pTZ-Δ*bccrh1*. To construct the *bccrh1* over-expression vector, the full-length *bccrh1* ORF fused to HA tag at the C-terminus was cloned into the pH2G vector^5^. To generate an enzyme inactive BcCrh1 protein, the catalytic residues were mutated (E120Q/D122H/E124Q) by site-directed PCR mutagenesis using template cDNA, and the amplification fragment was cloned into pJET plasmid to generate the pJET-MBcCrh1 vector. For generation of fungal strains that overexpress the MBcCrh1 in wild type and *bccrh1* deletion background, the M*bccrh1* clone was introduced into pNAN-OGG (contains a nourseothricin resistance cassette)^47^, between the *Aspergillus nidulans* PoliC promoter (for over expression) or the *bccrh1* promoter, and *B. cinerea* Tgluc terminator. Mutagenesis of the conserved cysteine residues at the N′ end (C26A/C33A/E120Q/D122H/E124Q) were generated by site-directed PCR mutagenesis and the amplified fragments were cloned into pNAN-OGG vector. For protein expression and purification, the sequence encoding mature proteins without the signal peptide was cloned into pET-14b (+) (Novagen) to generate the expression vector pET14b-6xHis-BcCrh1. Expression vectors of truncated-BcCrh1 were constructed in a similar way. To generate the constructs used for yeast-two hybrid assay (Y2H), the ORFs of target genes without the secretion sequence were amplified using PCR or site-directed mutagenesis PCR (C26A, C33A, C26A/C33A/ and Δ21-38), and the PCR amplification products were cloned into the pGADT7 and pGBKT7 yeast two hybrid vectors.

### DNA and RNA analyses

Genomic DNA and total RNA extractions were performed as described previously^5^. For cDNA synthesis, total RNA was treated with DNase I (Thermo Scientific) and then the first-strand cDNA was synthesized from 1 μg of DNA-free RNA using the RevertAid First Strand cDNA Synthesis Kit (Thermo Scientific). qRT-PCR was performed with SYBR Premix Ex Taq II (Takara, Dalian, China) using Mx3000P (Stratagene, La Jolla, CA, USA) or StepOne (Applied Biosystems) Real-time PCR instruments. Relative fold change of mRNA levels was normalized with the *B. cinerea bcgpdh* (BC1G_05277) or bean *actin-11* (GenBank accession no. EH040443.1) genes.

### *A. tumefaciens*-mediated transient expression

Constructs were introduced into *A. tumefaciens* strain GV3101 by electroporation, and Agrobacterium infiltration assays was performed on *N. benthamiana* plants that were grown for 4 to 5 weeks in the greenhouse as described previously^48^. Unless otherwise mentioned, leaves were photographed five days after infiltration. DAB staining of ROS was performed two days after agro-infiltration of leaves as previously described^49^.

### Leaf infiltration assay with purified proteins

To test the cell death-inducing activity of recombinant proteins, 2.75 - 55 μM protein solutions were infiltrated into leaves of *N. benthamiana*, and leaves of *S. lycopersicum*, *A. thaliana* and *Z. mays* were infiltrated with 11 μM protein solution. Infiltrated leaves were photographed 2-3 days after infiltration. To test the induced plant defense responses, total RNA was extracted from *S. lycopersicum* leaves at 0 h, 24 h and 48 h after infiltration with recombinant protein solutions (11 μM) and qRT-PCR analysis was performed for measurement of plant defense-related gene expression levels.

### Uptake assay of GFP-labeled proteins by plant cells

Tomato cell cultures (MsK8) were incubated for 30 min with incubation buffer (10 mM phosphate buffer, pH 7, and 0.1% BSA) followed by the addition of GFP-tagged peptide solutions (5.5 μM) for 18 h. The cells were subsequently washed three times in incubation buffer and visualized by confocal laser microscopy.

### Protein analysis

Plasmids used for proteins expression were transformed into *E. coli* strain BL21 (DE3). Expression and purification of the recombinant proteins were performed using Ni-NTAresin (GEHealthcare) according to previously described protocol^50^. For analysis of proteins following agroinfiltration, proteins were extracted from plant samples as previously described^5^. Immunoblots were performed with anti-GFP tag primary monoclonal antibody (Sigma-Aldrich) as described by^50^.

### Fluorescence and confocal microscopy

Samples were harvested from *N. benthamiana* leaves two days after the leaves were infiltrated with agrobacterium. For plasmolysis, samples were infiltrated with 0.8 M mannitol solution 20 min. For subcellular localization of GFP-labeled BcCrh1 in fungal cells during saprophytic growth, conidia were suspended in PDB and 10μl droplets of spore suspension containing 1×10^4^ conidia ml^−1^ were placed on a glass slides and incubated at 20°C. Confocal microscopy was performed with a Zeiss LSM780 confocal microscope system as described previously^51^. Epifluorescence and light microscopy were performed with a Zeiss Axio imager M1 microscope. Differential interference microscopy (DIC) was used for bright field images. DAPI filter (340-390 and 420-470 nm excitation and emission, respectively) was used for visualization of DAPI-stained nuclei and CMAC stained vacuoles. eGFP and mCherry fluorescence were collected by using an excitation laser wavelengths of 488 nm and 561 nm, respectively. For GFP, emission was collected in the range of 493-535 nm. Images were captured with a Zeiss AxioCam MRm camera.

### Transformation of *B. cinerea* and characterization of the transformants

PEG-Mediated protoplast transformation of *B. cinerea* was performed according to published protocols^52^. Transgenic strains are listed in Supplementary Table S2. At least three independent single spore isolates from independent colonies were obtained for each strain. The ΔBcCrh1 colonies were confirmed by PCR to verify deletion of the *bccrh1* gene, the level of *bccrh1* gene expression in overexpression strains was determined by qRT-PCR. The phenotypic assays, including mycelial growth rates, conidiation, conidial germination and stress tolerance were performed as previous description^52, 53^.

### Pathogenicity and infection-related assays

Pathogenicity assay with *B. cinerea* was performed as described previously^52^. For bean plants, 10-day-old primary leaves were inoculated with 7.5 μl of conidia suspension containing 2 ×10^5^ conidia ml^−1^. For *A. thaliana* plants, leaves of 4 to 5 weeks-old plants were similarly inoculated with 5 μl conidial suspension. Infection intensity was evaluated by measurement of lesion diameter 72 hpi. For infection cushion formation assay, conidia were suspended in GB5+2% Glucose, 10 μl were placed on glass slides and incubated in a moistened chamber at 20°C for 36 h. For localization of GFP-labeled BcCrh1 during infection, conidia were suspended in PDB medium and 10μl droplets of spore suspension containing 1×10^4^ conidia ml^−1^ were placed on onion epidermal cells and incubated at 20°C with a moistened condition. Samples were prepared for microscopy observation at designated time points.

### Staining dyes and procedures

Cotton blue and trypan blue staining of fugal hyphae for penetration assay of plant leaves were performed according to published procedures^54^. DAB staining of ROS was conducted based on the published description^55^. Viability nuclei staining with DAPI was performed as previously described^56^. For the staining of vacuoles with CMAC, cover glass cultures were first incubated in GB medium containing 10 μM CMAC for 20 min, followed by washing with fresh dye-free medium.

### Transformation of *A.thaliana*

*Agrobacterium*-mediated transformation of *A.thaliana* flowers was performed using the floral dip method according to a published protocol^57^. Three independent transgenic lines were generated with *bccrh1* over expression and empty vector (control) plasmids. Gene and protein expression were confirmed by PCR and western blot analysis, respectively. Homozygous T3 seeds were selected and used for all experiments.

### Yeast transformation and yeast two-hybrid assay

The Matchmaker Gold Yeast Two Hybrid (Y2H) System (Clontech, Palo Alto, CA, USA) was used to verify proteins interaction. The bait (pGBKT7) and prey (pGADT7) plasmids were co transformed into yeast strain AH109 using the LiAc/SS carrier DNA/PEG method^58^. Yeast transformants were screened on selective dropout (SD)/ Trp Leu medium to select yeast cells containing the desired plasmids (pGADT7 and pGBKT7). Positive protein interactions were assessed on SD/ Trp Leu His Ade medium supplemented with X-α-galactosidase (X-α-gal) after incubated at 28 °C for four days.

## Acknowledgments

We are grateful to Prof. Adi Avni and Mrs. Orian Taig for providing and assistance with tomato cell cultures. This work was supported by a BARD IS-4937-16

---

**Supplementary figure 1.**
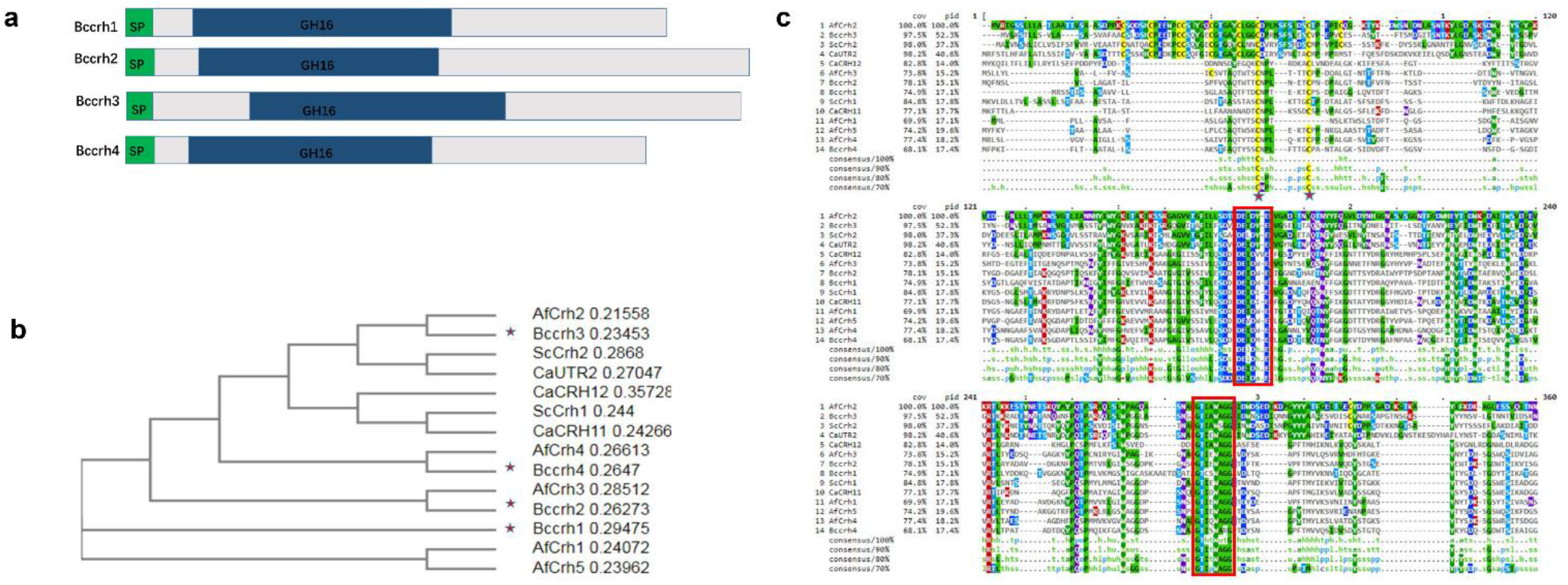
Phylogenetics and multiple sequence alignment of *B. cinerea* Crh family proteins. **a**, Schematic presentation of predicted signal peptide (green) and GH16 domain (blue) of BcCrh proteins. Multiple sequence alignment (MSA) analysis of *B. cinerea* Crh family proteins and their homologues from other species was generated by Clustal Omega online tool. Due to the C-terminus residues with low sequence identity, 1-277 of BcCrh1 was used for alignment. **b**, Neighbour-joining tree was generated based on the MSA result. The *B. cinerea* proteins are marked by stars. **c**, MSA results reformatted by Mview. The highly conserved Cys residues in all Crh family homologues from different species are marked with red stars. Other conserved residues are shaded by colors. The conserved catalytic activity domain and the acceptor sugar binding domain are boxed by red rectangle. Af – *Aspergillus fumigatus*, Sc – *Saccharomyces cerevisiae*, Ca – *Candida albincas*.

**Supplementary figure 2.**
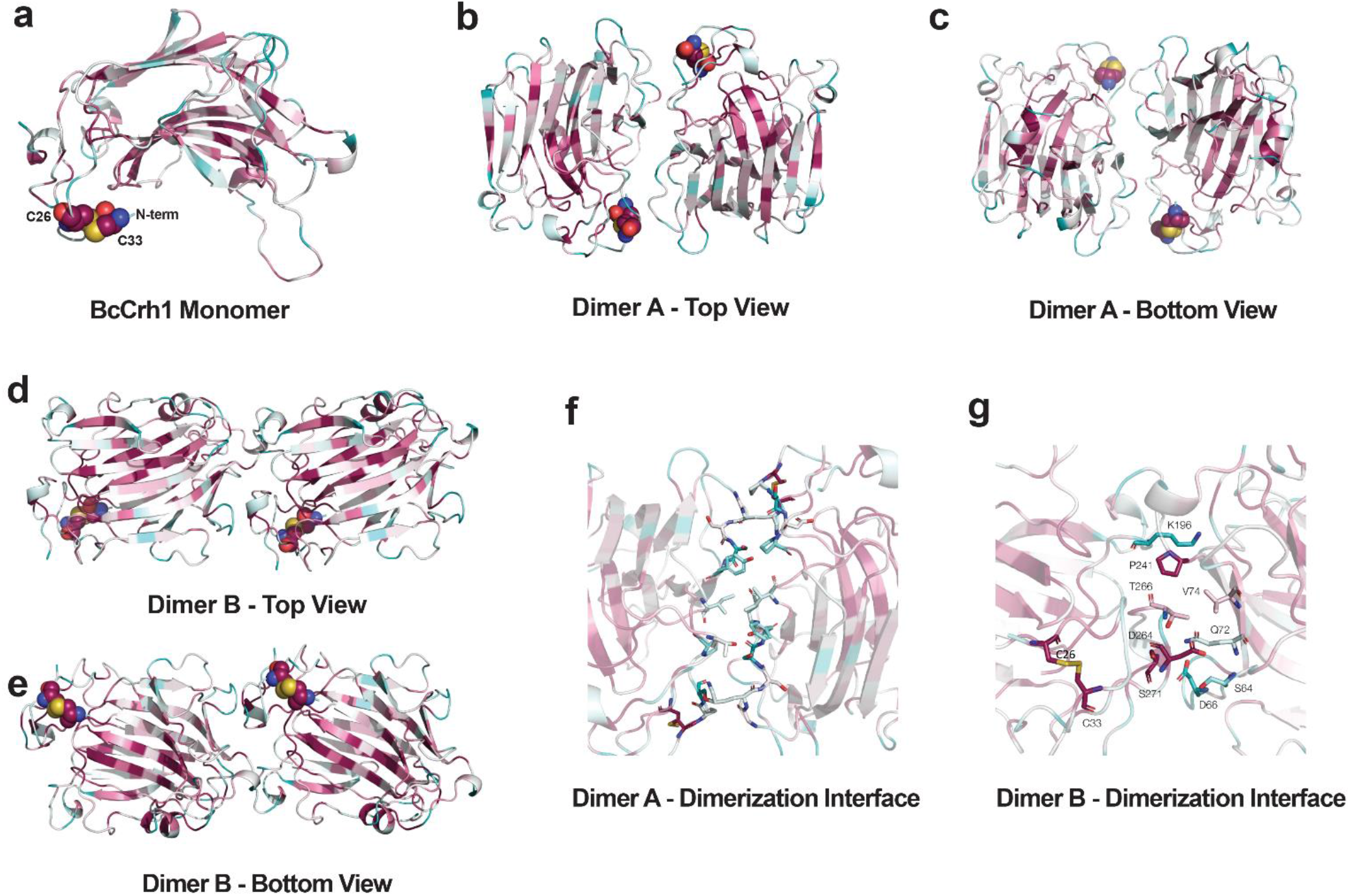
Predicted 3D structural models of BcCrh1. HHpred was used for modeling template search against PDB database. The AfCrh5 protein (PDB entry 6IBW, UniProt entry Q8J0P4) showed 48% sequence identity to BcCrh1 (Probability 100%, E-value 1.1e-32) and was used as a template for modeling. ConSurf was used to collect homologues from the clean UniProt database, the pairwise alignment of B1084 (later called BcCrh1) and 6IBW was deduced from the multiple sequence alignment. Different models were generated by MODELLER, and finally the model result with the predicted lowest energy was chosen. **a**, Overall structure of BcCrh1 monomer model. The C26 and C33 at the N-terminus highlighted in colored-spheres form a disulfide bond and are both highly conserved. **b-g**, Predicted potential dimerization interfaces of BcCrh1. PISA and the crystal structure of 6IBW were used to test potential dimerization interfaces of BcCrh1. Two different topologies of the dimer were predicted based on the analysis. Schematic ribbon diagram of dimerized BcCrh1 model are shown (the top view, bottom view and dimerization interface). (**b**, **c and f**) “head-to-head” topology of the dimer. (**d**, **e and g**) “head-to-tail” topology of the dimer.

**Supplementary figure 3.**
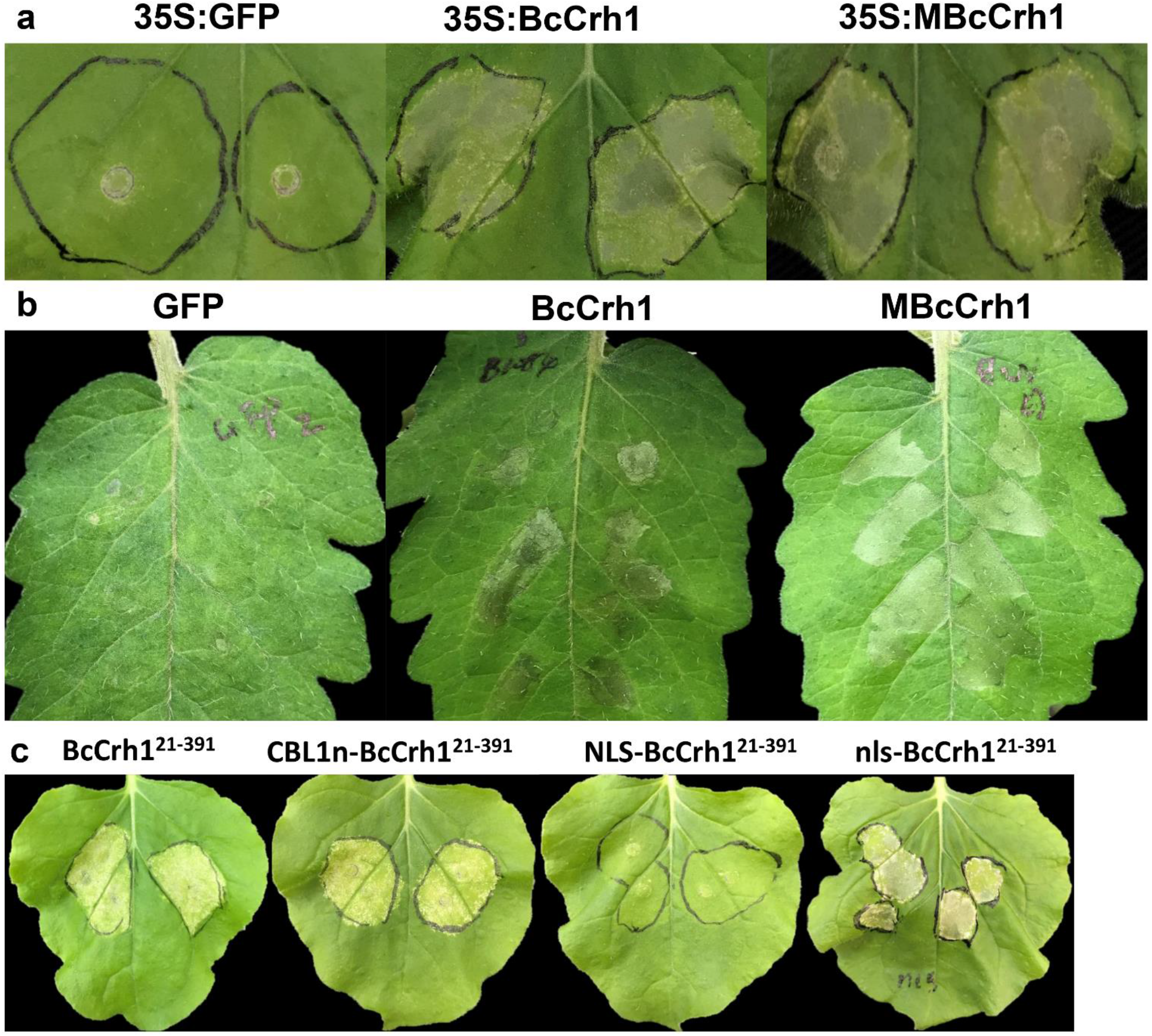
Induction of plant cell death by BcCrh1. **a**, *N. benthamiana* leaves were agroinfiltrated with GFP (35S:GFP), and native (35S:BcCrh1) or the enzymatic inactive (35S:MBcCrh1) BcCrh1. Images were taken five days after agroinfiltration. **b**, Tomato leaves were infiltrated with 11 μM of purified BcCrh1, MBcCrh1 and GFP proteins. Images were taken two days after infiltration. **c**, Representative *N. benthamiana* leaves five days after agroinfiltration with constructs expressing BcCrh1^21-391^, CBL1n-BcCrh1^21-391^, NLS-BcCrh1^21-391^ and nls-BcCrh1^21-391^.

**Supplementary figure 4.**
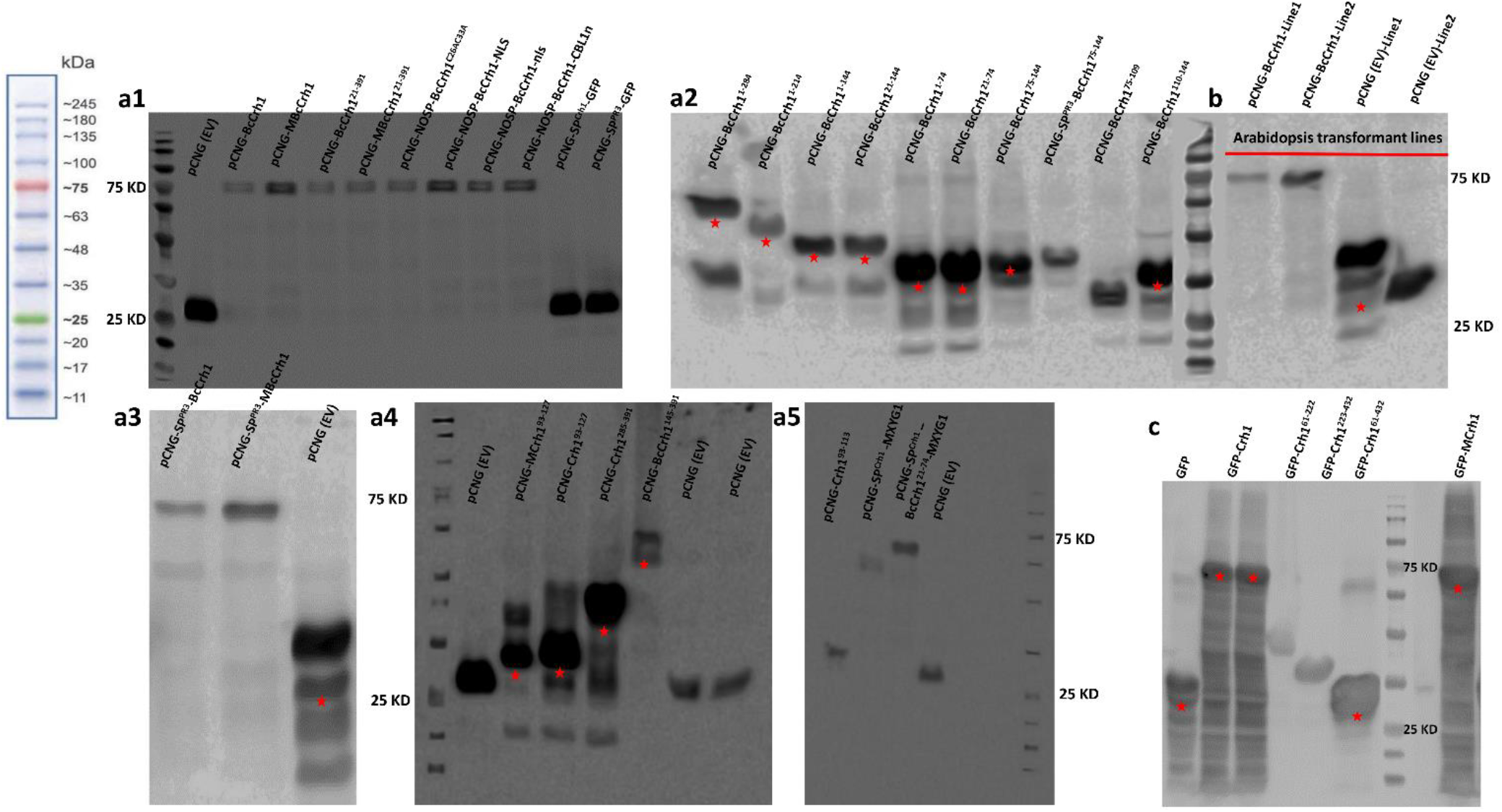
Detection of recombinant proteins by western blot. *α*-GFP antibody was used to detect expression of the indicated prtoeins. **a1-a5**, Immunoblot analysis of protein extracts from *N. benthamiana* leaves transiently expressing the different proteins fused to GFP. **b**, Immunoblot analysis of proteins isolated from *A. thaliana* transgenic lines expressing the BcCrh1-GFP (Line1 and Line2) and empty vector (EV). **c**, Immunoblot analysis of the indicated proteins fused with GFP tag from *E. coli*.

**Supplementary figure 5.**
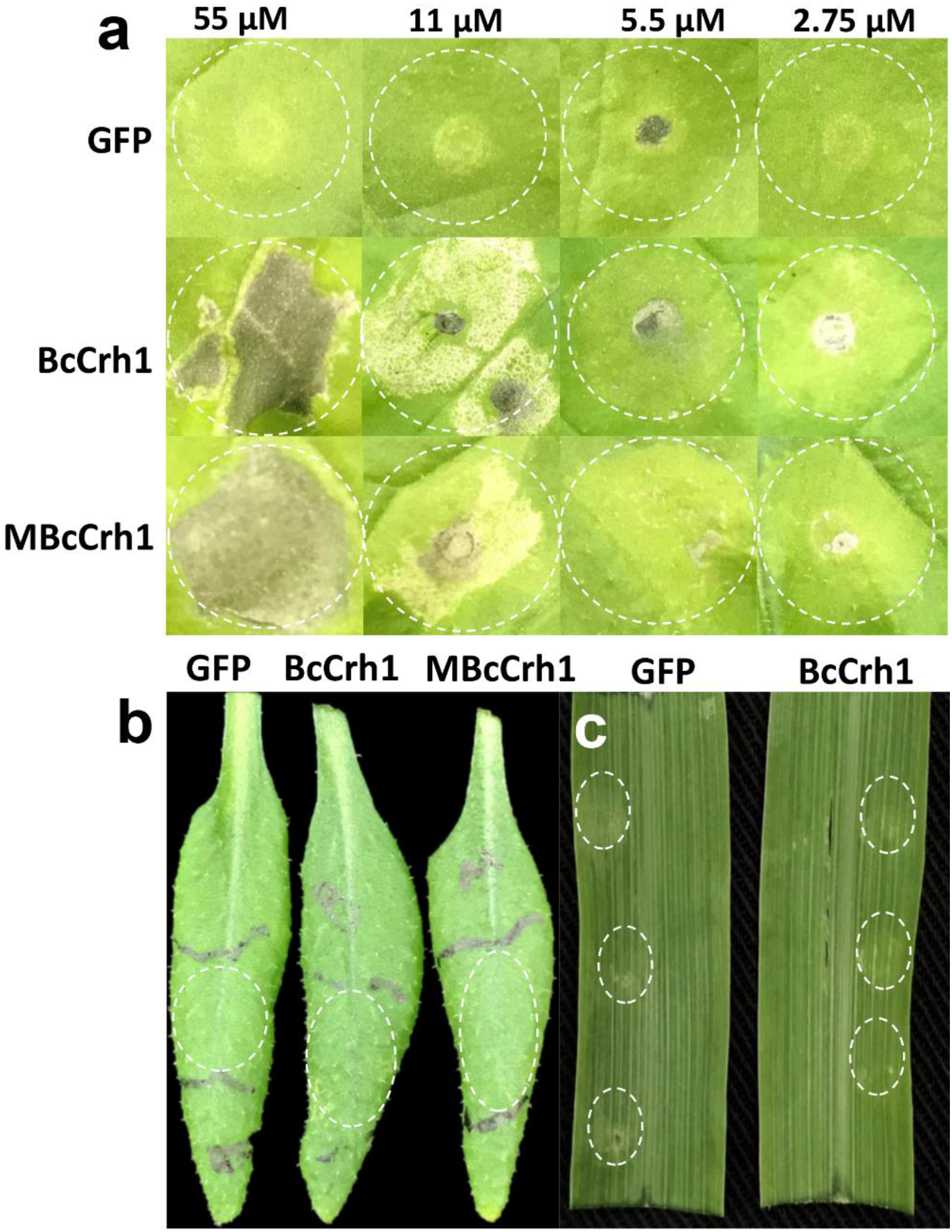
Infiltration of leaves with purified BcCrh1 protein. **a**, Response of *N. benthamiana* leaves to different concentrations of purified BcCrh1 and MBcCrh1 protein. Pictures were taken two days after treatment. **b, c**, Cell death response assay on *A. thaliana* (**b**) and maize (*Zea mays*) (**c**) leaves. Leaves were infiltrated with 11 μM of purified protein, pictures of representative leaves were taken two days after treatment. White dash line shows the boundary of infiltrated leaves.

**Supplementary figure 6.**
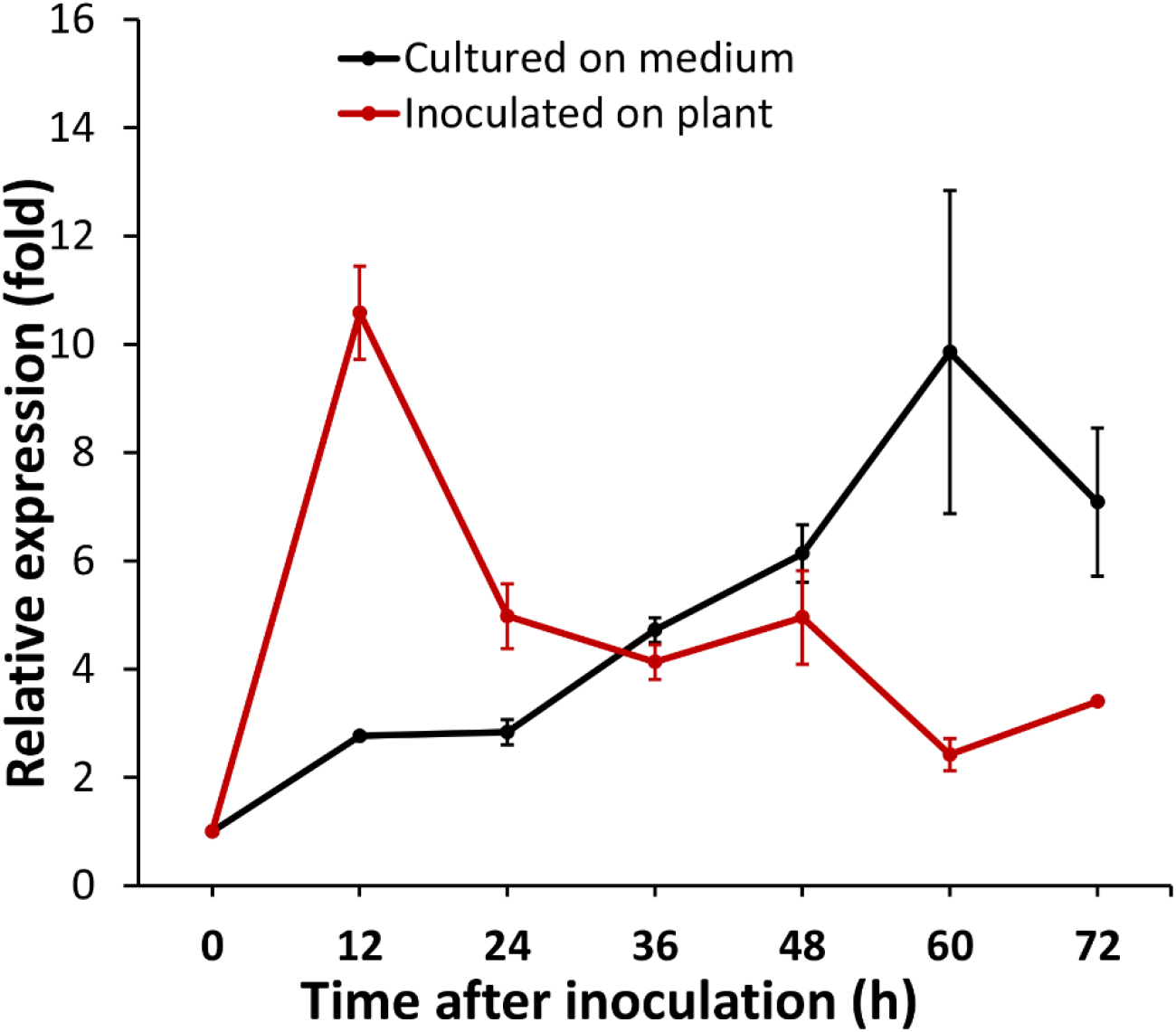
Transcript analysis of *bccrh1* during saprophytic and pathogenic fungal development. *bccrh1* expression levels were evaluated by qRT-PCR with the *bcgpdh* as a reference gene for normalization. Mean and standard deviations were calculated from three biological replicates. The transcript relative levels were calculated using the comparative Ct method, and the expression level of *bcCrh1* inoculated on plants (black line) or in Gamborg’s B5 medium (red line) at 0 h was set as 1.

**Supplementary figure 7.**
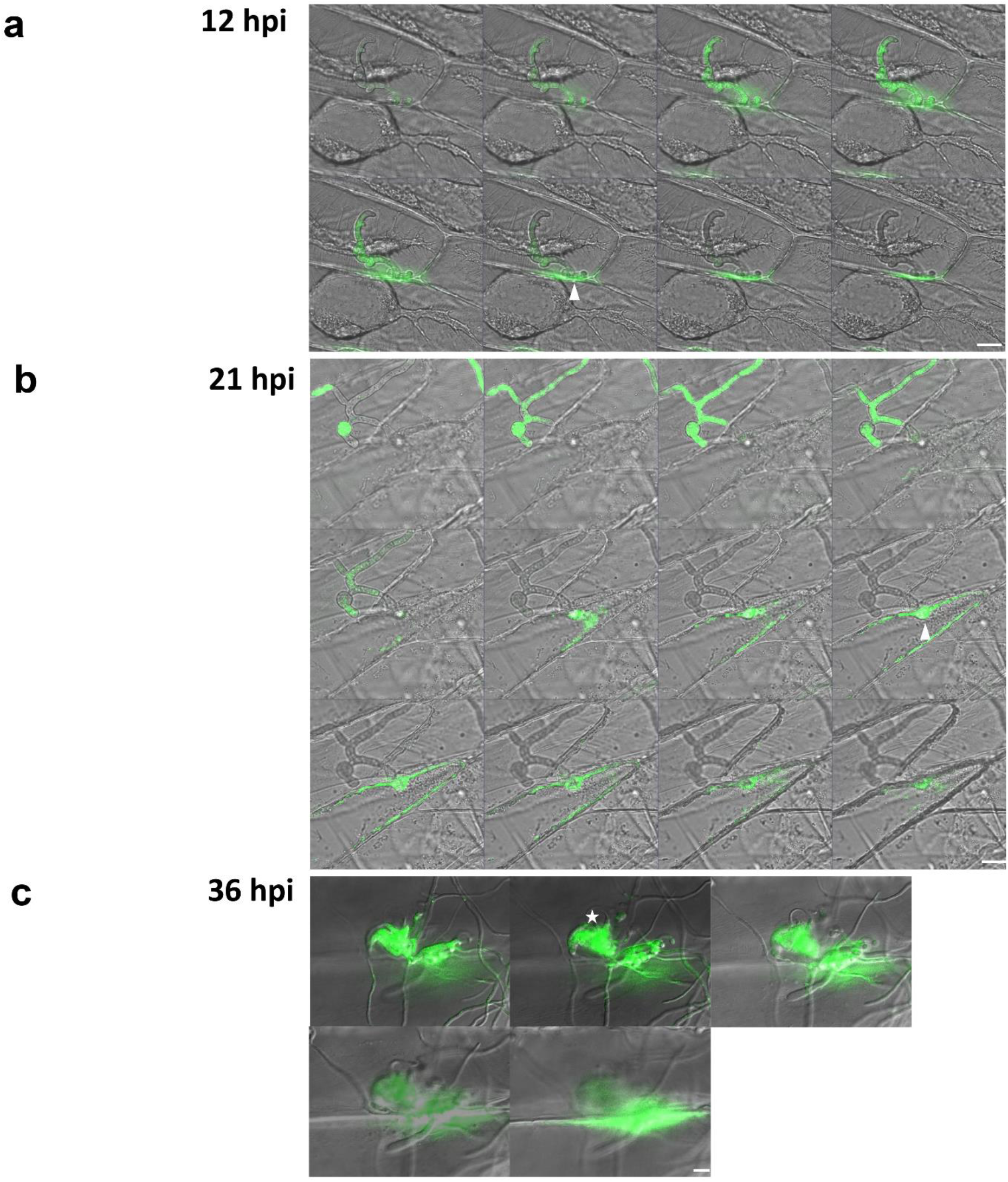
Secretion of Bccrh1 during plant infection. Onion epidermis was inoculated with spore suspension of a *B. cinerea* strain that expresses BcCrh1-GFP fusion protein. Samples were scanned by a confocal microscope. Images show continuous z-series of different infection states. **a, b**, Protein localization to hyphal tips 12 and 21 hpi (arrowhead). **c,** Protein localization in infection structures and in the plant apoplast (36 hpi, asterisks). Bar = 20 μm.

**Supplementary figure 8.**
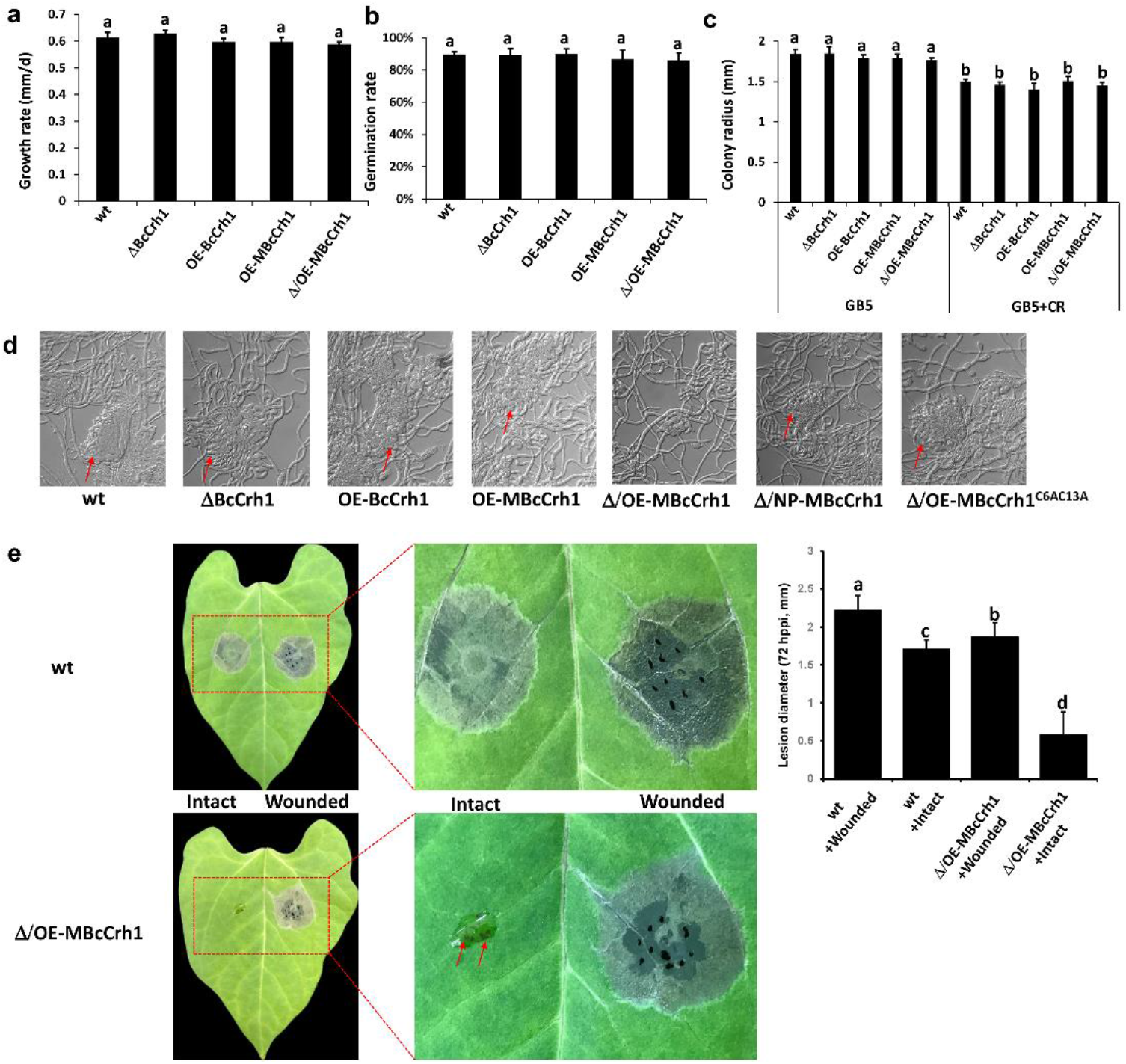
Developmental phenotypes of *B. cinerea* mutant strains. **a**, Hyphal growth rate. Fungi were grown on GB5+Glucose plates at 22°C with continuous fluorescent light. Radial growth was measured every day for three days and growth rate (mm/d) was calculated. Data represent the means and standard deviations from three independent biological replicates. **b**, Conidia germination. Values stand for the means and standard deviations from three independent biological replicates, at least 120 conidia were counted in each replication. **c**, Effect of cell wall stressor on colony radial growth. Fungi were inoculated on GB5+Glucose medium with or without 300 μg/ml congo red (CR) and colony diameter was measured after three days. Data represent the means and standard deviations from three independent biological replicates. Same letters in the graphs indicate no statistical differences at *P* ≤ 0.01 according to one-way ANOVA. **d**, Infection cushion formation and morphology. Conidia were suspended in GB5+2% Gluc medium, density was adjusted 10^4^ conidia ml^−1^ and 10 μl of the suspension were applied to a glass slide, and the slides were incubated in a moist chamber at 22°C. Images show samples after 36 h of incubation. Arrow indicates a typical infection cushion. Notably, at this stage, all strains except Δ/OE-MBcCrh1 formed normal infection cushions. Strain designation: wt – B05.10 wild type, ΔBcCrh1 – *bccrh1* deletion, OE-BcCrh1 – *bccrh1* over-expression, OE-MBcCrh1 – over-expression of enzyme inactive *bccrh1*, Δ/NP-MBcCrh1 – expression of the enzyme inactive *bccrh1* in background of *bccrh1* deletion, Δ/OE-MBcCrh1C6AC13A - over-expression of enzyme inactive *bccrh1* with mutations in C6 and C13 in background of *bccrh1* deletion. **e,** Bean leaves (Intact and wounded) were inoculated with spore suspensions of *B. cinerea* wild type (wt) and Δ/OE-MBcCrh1. At 3 dpi, typical lesions were present and measured. Data collected from three independent biological replicates. Different letters indicate statistical differences at *P* ≤ 0.01 using one-way ANOVA.

**Supplementary figure 9.**
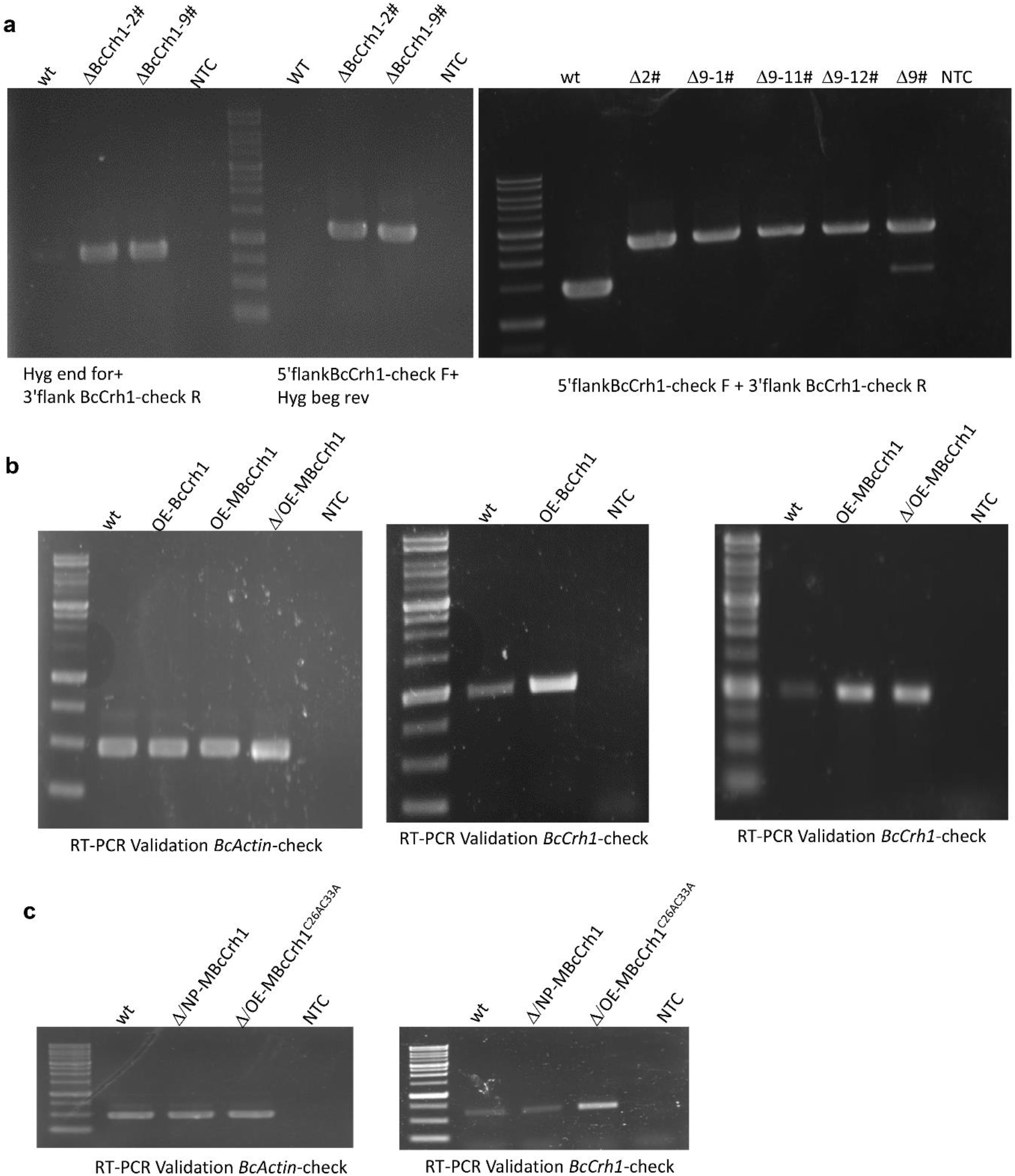
PCR validation of *B. cinerea* mutant strains. **a**, Analysis of *bccrh1* deletion strains DBcCrh1-2 and DBcCrh1-9. **b** - **c**, RT-PCR analysis of *bccrh1* expression in different over-expression strains. The *bcactin* gene was used to normalize different samples. NTC – no template control, ddH_2_O was used as negative control.

**Supplemental Table 1.**
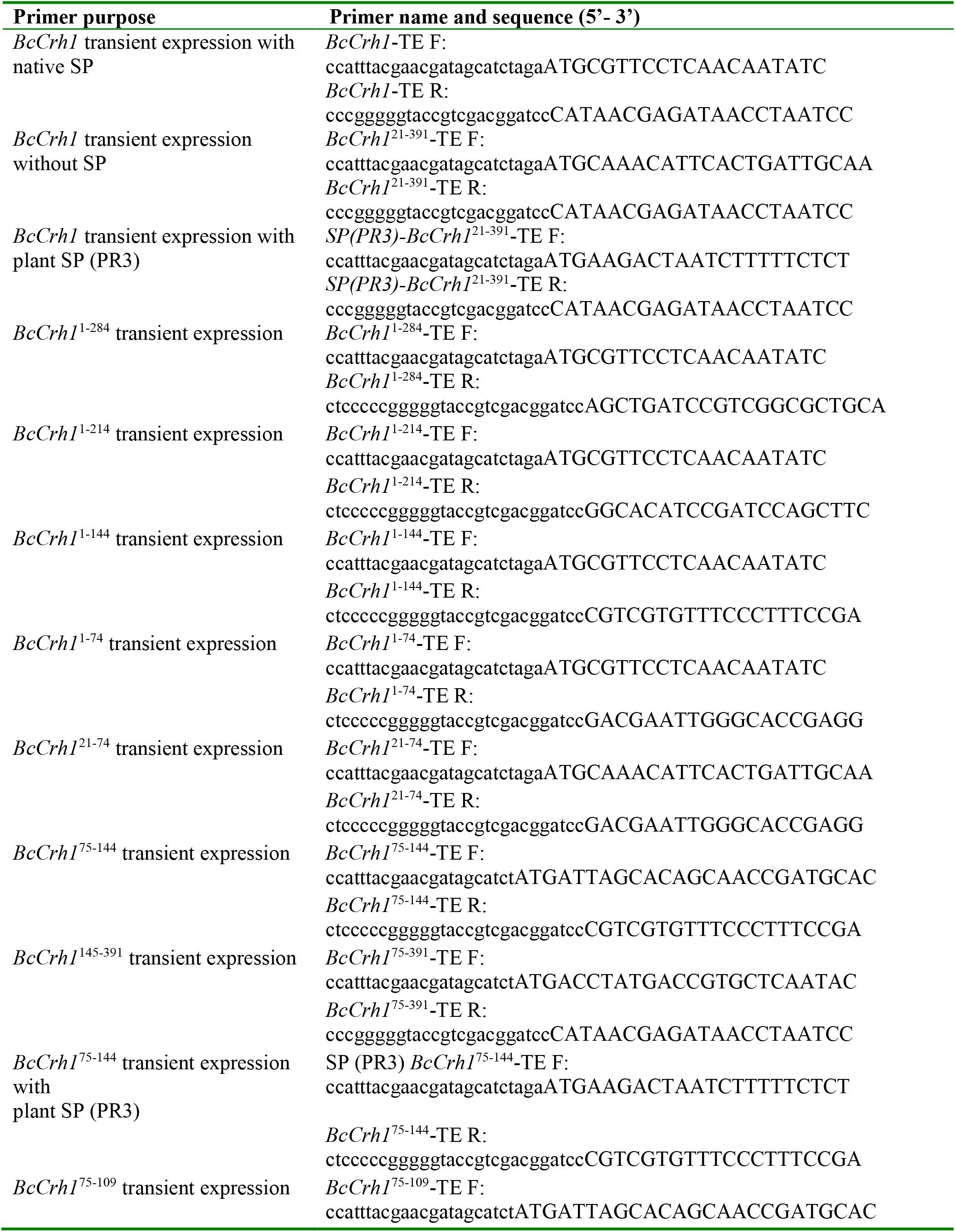
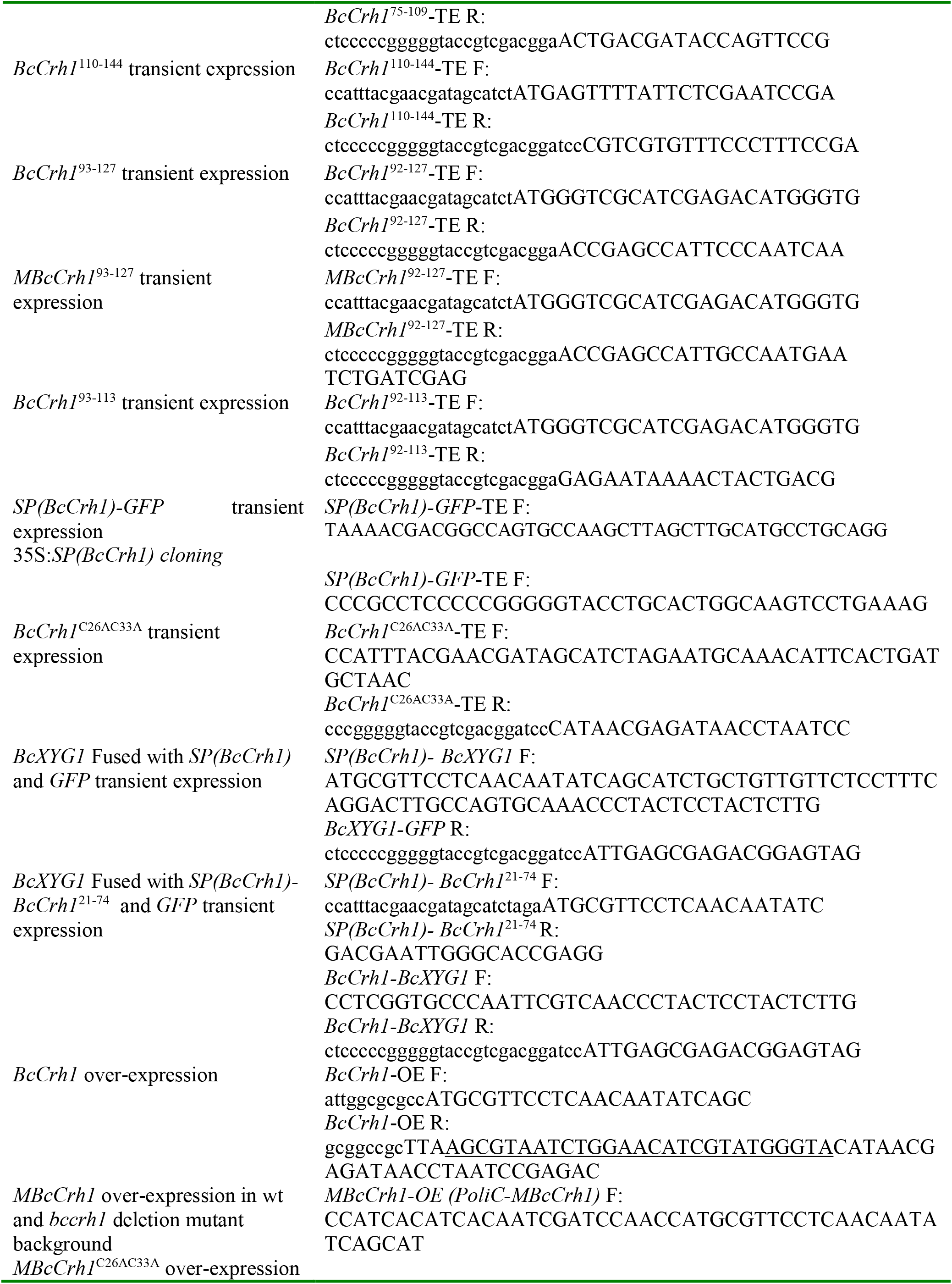
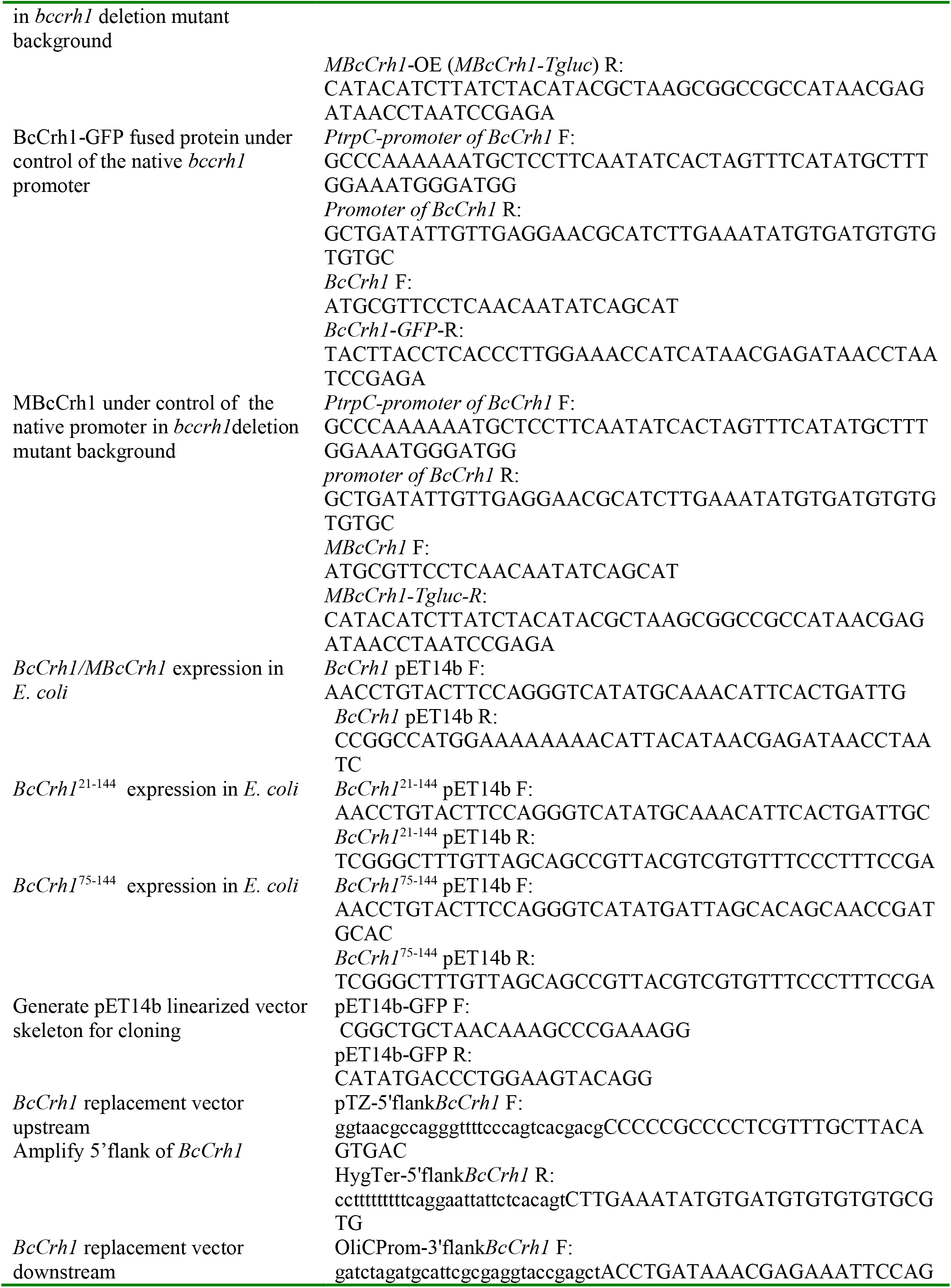
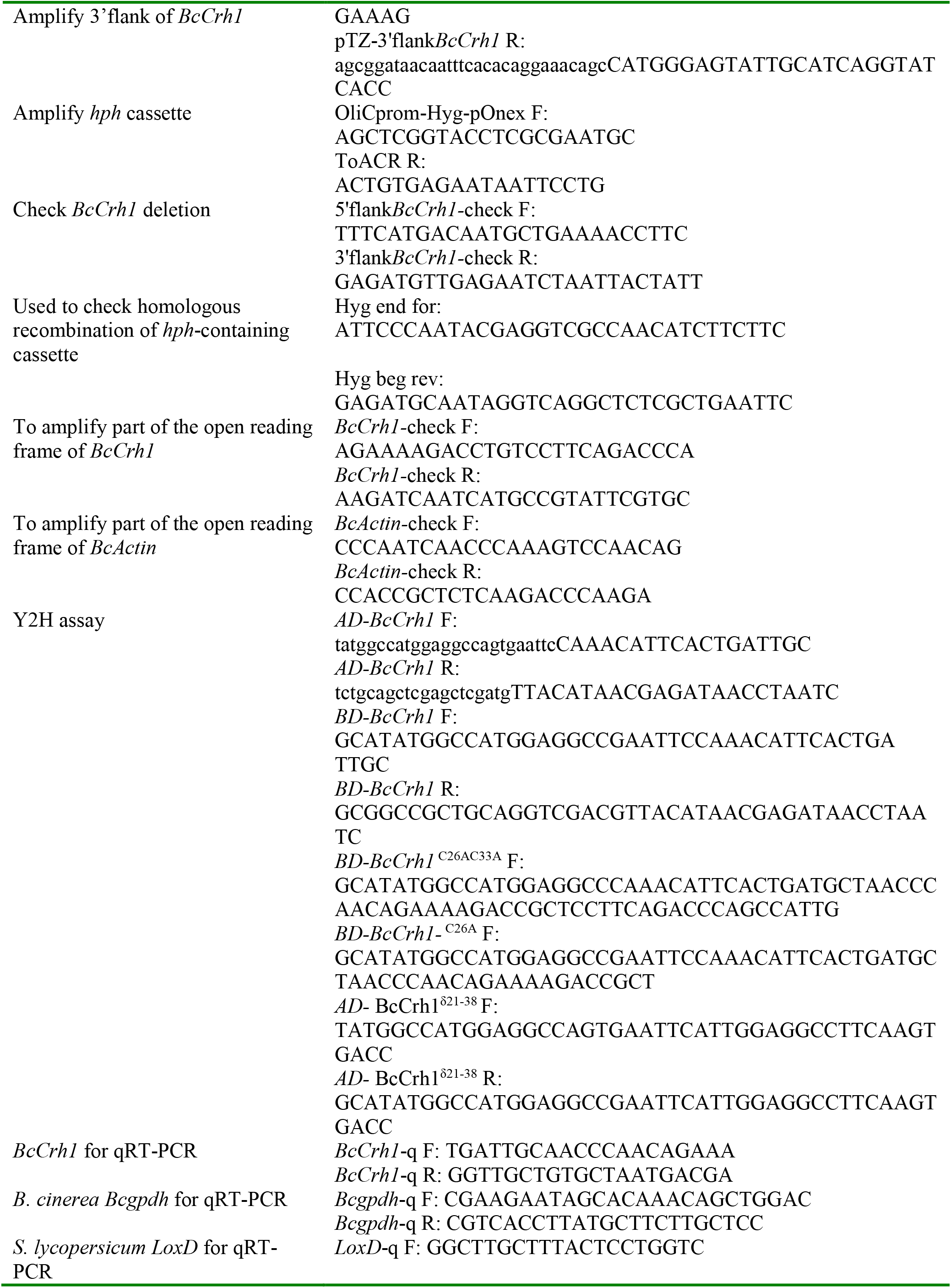
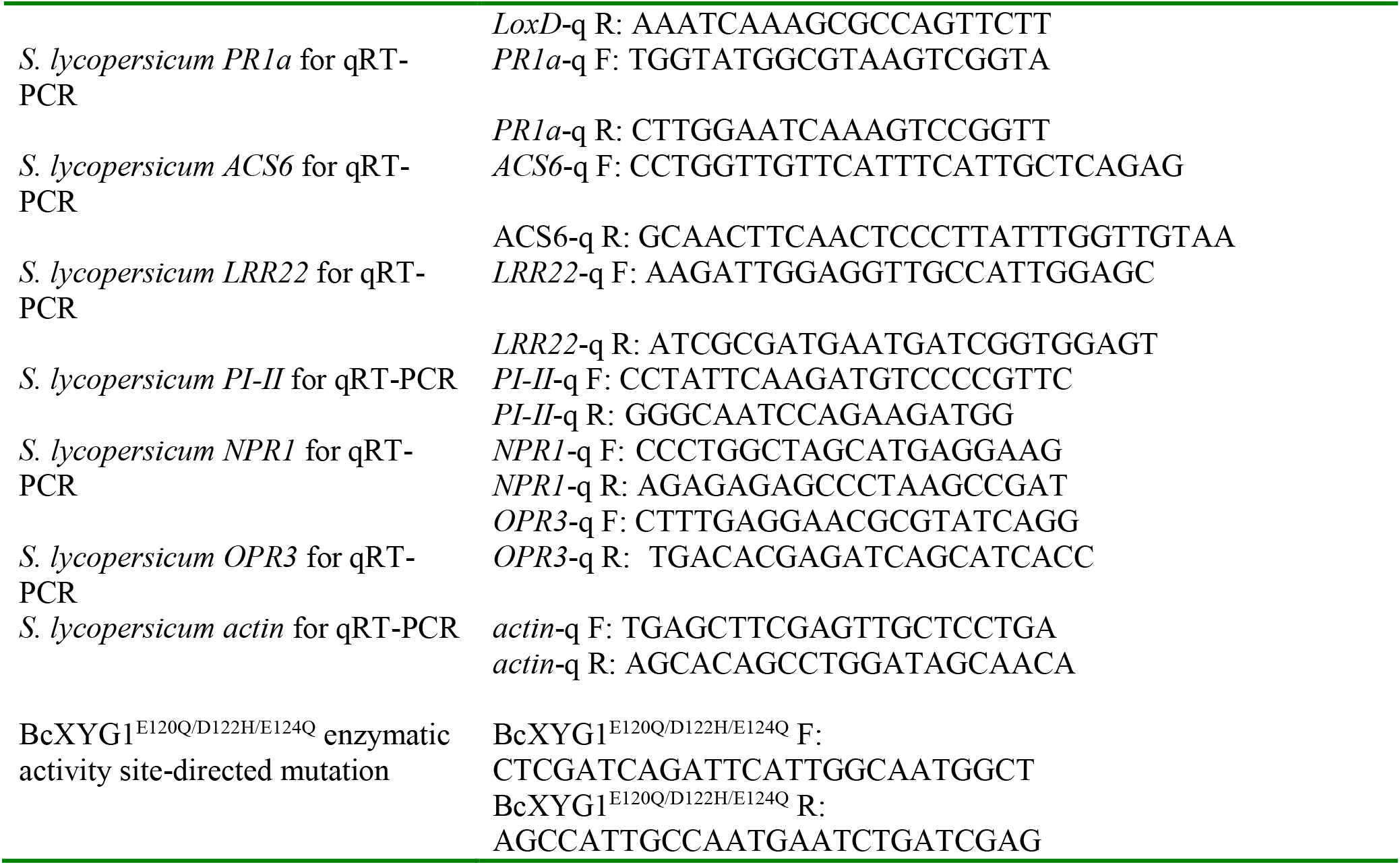
List of Oligonucleotides used in this study.

**Supplemental Table 2.** List of *B. cinerea* strains used in this study.

| Strain | Genotype and description | Source |
| --- | --- | --- |
| B05.10 | Haploidized *B. cinerea* wild type strain | Shlezinger et al., (2016) |
| $\Delta bccrh1$ | *bccrh1* deletion mutant | This work |
| OE-BcCrh1 | Over expression strain of the native form of BcCrh1 | This work |
| OE-MBcCrh1 | Over expression strain of the mutated form of BcCrh1 that has no enzymatic activity in wild-type background | This work |
| NP-BcCrh1-GFP | BcCrh1-GFP fused protein under control of the native *bccrh1* promoter | This work |
| $\Delta$/OE-MBcCrh1 | Over expression strain of the mutated form of BcCrh1 that has no enzymatic activity in *bccrh1* deletion background | This work |
| $\Delta$/NP-MBcCrh1 | Mutant strain of the enzymatic inactive BcCrh1 protein under control of the native *bccrh1* promoter in *bccrh1* deletion background | This work |
| $\Delta$/OE-MBcCrh1<sup>C26AC33A</sup> | Over expression strain of the mutated form of BcCrh1 that lost both enzymatic activity and homodimer activity in *bccrh1* deletion background | This work |

## Notes

### Competing Interest Statement

The authors have declared no competing interest.

## References

1. Amselem J, et al. Genomic analysis of the necrotrophic fungal pathogens Sclerotinia sclerotiorum and Botrytis cinerea. PLoS genetics 7, (2011).

2. Dean R, et al. The Top 10 fungal pathogens in molecular plant pathology. Molecular plant pathology 13, 414–430 (2012).

3. Eizner E, Ronen M, Gur Y, Gavish A, Zhu W, Sharon A. Characterization of Botrytis–plant interactions using PathTrack©—an automated system for dynamic analysis of disease development. Molecular plant pathology 18, 503–512 (2017).

4. Shlezinger N, et al. Anti-apoptotic machinery protects the necrotrophic fungus Botrytis cinerea from host-induced apoptotic-like cell death during plant infection. PLoS pathogens 7, e1002185 (2011).

5. Zhu W, et al. BcXYG1, a secreted xyloglucanase from Botrytis cinerea, triggers both cell death and plant immune responses. Plant physiology 175, 438–456 (2017).

6. Bailey BA. Purification of a protein from culture filtrates of Fusarium oxysporum that induces ethylene and necrosis in leaves of Erythroxylum coca. Phytopathology 85, 1250–1255 (1995).

7. Pemberton CL, Salmond GP. The Nep1 like proteins—a growing family of microbial elicitors of plant necrosis. Molecular Plant Pathology 5, 353–359 (2004).

8. Lenarčič T, et al. Eudicot plant-specific sphingolipids determine host selectivity of microbial NLP cytolysins. Science 358, 1431–1434 (2017).

9. Lenarčič T, et al. Molecular basis for functional diversity among microbial Nep1-like proteins. PLoS pathogens 15, e1007951 (2019).

10. Seidl MF, Van den Ackerveken G. Activity and phylogenetics of the broadly occurring family of microbial Nep1-like proteins. Annual review of phytopathology 57, 367–386 (2019).

11. Have At, Mulder W, Visser J, van Kan JA. The endopolygalacturonase gene Bcpg1 is required for full virulence of Botrytis cinerea. Molecular Plant-Microbe Interactions 11, 1009–1016 (1998).

12. Noda J, Brito N, González C. The Botrytis cinerea xylanase Xyn11A contributes to virulence with its necrotizing activity, not with its catalytic activity. BMC plant biology 10, 38 (2010).

13. Yang Y, Yang X, Dong Y, Qiu D. The Botrytis cinerea xylanase BcXyl1 modulates plant immunity. Frontiers in microbiology 9, 2535 (2018).

14. Zhang Y, Zhang Y, Qiu D, Zeng H, Guo L, Yang X. BcGs1, a glycoprotein from Botrytis cinerea, elicits defence response and improves disease resistance in host plants. Biochemical and biophysical research communications 457, 627–634 (2015).

15. Frías M, González MA, González C, Brito N. A 25-residue peptide from Botrytis cinerea xylanase BcXyn11A elicits plant defenses. Frontiers in plant science 10, 474 (2019).

16. Pérez Hernández A, González M, González C, Brito N. The elicitor protein BcIEB1 and the derived peptide ieb35 provide long term plant protection. Plant Pathology 69, 807–817 (2020).

17. Kars I, Krooshof GH, Wagemakers L, Joosten R, Benen JA, Van Kan JA. Necrotizing activity of five Botrytis cinerea endopolygalacturonases produced in Pichia pastoris. The Plant Journal 43, 213–225 (2005).

18. Rodríguez-Peña JM, Cid VJ, Arroyo J, Nombela C. A novel family of cell wall-related proteins regulated differently during the yeast life cycle. Molecular and cellular biology 20, 3245–3255 (2000).

19. Pardini G, et al. The CRH family coding for cell wall glycosylphosphatidylinositol proteins with a predicted transglycosidase domain affects cell wall organization and virulence of Candida albicans. Journal of Biological Chemistry 281, 40399–40411 (2006).

20. Fang W, et al. Mechanisms of redundancy and specificity of the Aspergillus fumigatus Crh transglycosylases. Nature communications 10, 1–10 (2019).

21. Blanco-Ulate B, Morales-Cruz A, Amrine KC, Labavitch JM, Powell AL, Cantu D. Genome-wide transcriptional profiling of Botrytis cinerea genes targeting plant cell walls during infections of different hosts. Frontiers in plant science 5, 435 (2014).

22. González-Fernández R, Aloria K, Valero-Galván J, Redondo I, Arizmendi JM, Jorrín-Novo JV. Proteomic analysis of mycelium and secretome of different Botrytis cinerea wild-type strains. Journal of proteomics 97, 195–221 (2014).

23. Müller N, et al. Investigations on VELVET regulatory mutants confirm the role of host tissue acidification and secretion of proteins in the pathogenesis of Botrytis cinerea. New Phytologist 219, 1062–1074 (2018).

24. Schouten A, Van Baarlen P, Van Kan JA. Phytotoxic Nep1 like proteins from the necrotrophic fungus Botrytis cinerea associate with membranes and the nucleus of plant cells. New Phytologist 177, 493–505 (2008).

25. Frías M, González C, Brito N. BcSpl1, a cerato platanin family protein, contributes to Botrytis cinerea virulence and elicits the hypersensitive response in the host. New Phytologist 192, 483–495 (2011).

26. Fillinger S, Elad Y. Botrytis: the fungus, the pathogen and its management in agricultural systems. Springer (2016).

27. Chagué V, et al. Ethylene sensing and gene activation in Botrytis cinerea: a missing link in ethylene regulation of fungus-plant interactions? Molecular plant-microbe interactions 19, 33–42 (2006).

28. Blanco N, et al. Structural and functional analysis of yeast Crh1 and Crh2 transglycosylases. The FEBS journal 282, 715–731 (2015).

29. Hwang JS, Seo DH, Kim JY. Soluble forms of YlCrh1p and YlCrh2p, cell wall proteins of Yarrowia lipolytica, have β 1, 3 glycosidase activity. Yeast 23, 803–812 (2006).

30. Agrawal P, et al. CPPsite 2.0: a repository of experimentally validated cell-penetrating peptides. Nucleic acids research 44, D1098–D1103 (2016).

31. Birch PR, Rehmany AP, Pritchard L, Kamoun S, Beynon JL. Trafficking arms: oomycete effectors enter host plant cells. Trends in microbiology 14, 8–11 (2006).

32. Whisson SC, et al. A translocation signal for delivery of oomycete effector proteins into host plant cells. Nature 450, 115–118 (2007).

33. Presti LL, Kahmann R. How filamentous plant pathogen effectors are translocated to host cells. Current opinion in plant biology 38, 19–24 (2017).

34. Rehmany AP, et al. Differential recognition of highly divergent downy mildew avirulence gene alleles by RPP1 resistance genes from two Arabidopsis lines. The Plant Cell 17, 1839–1850 (2005).

35. Torto TA, et al. EST mining and functional expression assays identify extracellular effector proteins from the plant pathogen Phytophthora. Genome research 13, 1675–1685 (2003).

36. Schornack S, et al. Ancient class of translocated oomycete effectors targets the host nucleus. Proceedings of the National Academy of Sciences 107, 17421–17426 (2010).

37. Kemen E, et al. Gene gain and loss during evolution of obligate parasitism in the white rust pathogen of Arabidopsis thaliana. PLoS biology 9, (2011).

38. Dou D, et al. RXLR-mediated entry of Phytophthora sojae effector Avr1b into soybean cells does not require pathogen-encoded machinery. The Plant Cell 20, 1930–1947 (2008).

39. Kale SD, et al. External lipid PI3P mediates entry of eukaryotic pathogen effectors into plant and animal host cells. Cell 142, 284–295 (2010).

40. Plett JM, et al. A secreted effector protein of Laccaria bicolor is required for symbiosis development. Current Biology 21, 1197–1203 (2011).

41. Gu B, et al. Rust secreted protein Ps87 is conserved in diverse fungal pathogens and contains a RXLR-like motif sufficient for translocation into plant cells. PLoS One 6, (2011).

42. Di X, Gomila J, Ma L, van den Burg HA, Takken FL. Uptake of the Fusarium effector Avr2 by tomato is not a cell autonomous event. Frontiers in plant science 7, 1915 (2016).

43. Armenteros JJA, et al. SignalP 5.0 improves signal peptide predictions using deep neural networks Nature biotechnology 37, 420–423 (2019).

44. Krogh A, Larsson B, Von Heijne G, Sonnhammer EL. Predicting transmembrane protein topology with a hidden Markov model: application to complete genomes. Journal of molecular biology 305, 567–580 (2001).

45. Letunic I, Bork P. 20 years of the SMART protein domain annotation resource. Nucleic acids research 46, D493–D496 (2018).

46. Kostylev M, Otwell AE, Richardson RE, Suzuki Y. Cloning should be simple: Escherichia coli DH5α-mediated assembly of multiple DNA fragments with short end homologies. PLoS One 10, e0137466 (2015).

47. Schumacher J. Tools for Botrytis cinerea: new expression vectors make the gray mold fungus more accessible to cell biology approaches. Fungal genetics and biology 49, 483–497 (2012).

48. Wang Q, et al. Transcriptional programming and functional interactions within the Phytophthora sojae RXLR effector repertoire. The Plant Cell 23, 2064–2086 (2011).

49. Li Q, et al. A phytophthora capsici effector targets ACD11 binding partners that regulate ROS-mediated defense response in arabidopsis. Molecular plant 12, 565–581 (2019).

50. Ma L, Salas O, Bowler K, Bar-Peled M, Sharon A. UDP-4-Keto-6-Deoxyglucose, a Transient Antifungal Metabolite, Weakens the Fungal Cell Wall Partly by Inhibition of UDP-Galactopyranose Mutase. mBio 8, e01559–01517 (2017).

51. Pizarro L, Leibman-Markus M, Schuster S, Bar M, Meltz T, Avni A. Tomato prenylated RAB acceptor protein 1 modulates trafficking and degradation of the pattern recognition receptor LeEIX2, affecting the innate immune response. Frontiers in plant science 9, 257 (2018).

52. Ma L, Salas O, Bowler K, Oren Young L, Bar Peled M, Sharon A. Genetic alteration of UDP rhamnose metabolism in Botrytis cinerea leads to the accumulation of UDP KDG that adversely affects development and pathogenicity. Molecular plant pathology 18, 263–275 (2017).

53. Liu JK, et al. The key gluconeogenic gene PCK1 is crucial for virulence of Botrytis cinerea via initiating its conidial germination and host penetration. Environmental microbiology 20, 1794–1814 (2018).

54. Soulié MC, et al. Botrytis cinerea virulence is drastically reduced after disruption of chitin synthase class III gene (Bcchs3a). Cellular microbiology 8, 1310–1321 (2006).

55. Liu Z, Friesen T. DAB staining and visualization of hydrogen peroxide in wheat leaves. Bioprotocol 2, e309 (2012).

56. Barhoom S, Sharon A. cAMP regulation of “pathogenic” and “saprophytic” fungal spore germination. Fungal Genetics and Biology 41, 317–326 (2004).

57. Zhang X, Henriques R, Lin S-S, Niu Q-W, Chua N-H. Agrobacterium-mediated transformation of Arabidopsis thaliana using the floral dip method. Nature protocols 1, 641 (2006).

58. Gietz RD, Woods RA. Yeast transformation by the LiAc/SS Carrier DNA/PEG method. In: Yeast Protocol). Springer (2006).

